# Deletion of epithelial HKDC1 decelerates cellular proliferation and impairs mitochondrial function of tumorous epithelial cells thereby protecting from intestinal carcinogenesis in mice

**DOI:** 10.1101/2024.11.15.623798

**Authors:** Lea Järke, Saskia Weber-Stiehl, Kensuke Shima, Karlis Arturs Moors, Jerome Genth, Fenja Amrei Schuran, Lena Best, Markus Tschurtschenthaler, Burkhardt Flemer, Andreas Tholey, Christoph Kaleta, Jan Rupp, Philip Rosenstiel, Felix Sommer

## Abstract

**Background:** A metabolic switch favoring glycolysis over aerobic oxidative phosphorylation, namely the “Warburg effect”, represents a hallmark of cancer cells. Hexokinases (HK) catalyze the first step of glycolysis, thereby regulating its rate. Dysregulated HKDC1 (HK domain containing 1) expression has been associated with various cancer types and blocking HKDC1 prevents disease progression for hepatic carcinoma T cell lymphoma and lung adenocarcinoma, but its implication for colorectal cancer (CRC) remained unknown. Here, we functionally investigated the role of HKDC1 for intestinal carcinogenesis.

**Methods:** First, we analyzed HKDC1 expression in the intestinal mucosa of healthy controls (HC) and CRC patients and in different tumor tissues using transcriptomic data from publicly available databases. We then generated HKDC1-deficient human and murine colonic epithelial cell lines as well as intestinal organoids and profiled their phenotypic functions. Next, we screened for proteins interacting with HKDC1 by immunoprecipitation. Finally, we generated tumor-bearing *Apc*^Min/+^ mice with a conditional deletion of HKDC1 in intestinal epithelial cells and also performed a xenograft mouse model to test the role of HKDC1 for intestinal carcinogenesis *in vivo*.

**Results:** HKDC1 was found to be overexpressed in tumor compared to normal tissue of CRC patients. *In vitro,* HKDC1-deficient human Caco-2 and murine CMT-93 cells displayed reduced proliferation, altered susceptibility to cell death induction, and disrupted mitochondrial functions, particularly mitochondrial respiration. These altered cancer hallmarks were then corroborated in HKDC1-deficient normal and tumor-derived *Apc*^Min/+^ intestinal organoids. Immunoprecipitation and mass spectometry proteomic analyses revealed interactions of HKDC1 with several mitochondria-related proteins. *In vivo*, two distinct mouse models demonstrated that epithelial deletion of HKDC1 protected from carcinogenesis. First, *Apc*^Min/+^-*Hkdc1*^ΔIEC^ mice showed mildly improved disease phenotypes in the colon accompanied with reduced numbers of Ki67-positive proliferating epithelial cells. Finally, HKDC1-deficient Caco-2 cells completely failed to form any tumor mass in a xenograft model when implanted into immunodeficient mice.

**Conclusions:** We demonstrate that HKDC1 influences cancer cell proliferation and susceptibility to cell death, potentially through interactions with mitochondrial proteins that regulate membrane permeability, ultimately impacting intestinal carcinogenesis. Collectively, these findings highlight the significance of HKDC1 for CRC pathobiology, presenting it as a promising target for further investigation and potential therapeutic interventions.

## Introduction

Colorectal cancer (CRC) is among the leading causes of death worldwide and the third most prevalent cancer type (Siegel, Miller, et al., 2023; Siegel, Wagle, et al., 2023). Cancer cells display certain altered cellular characteristics termed hallmarks of cancer cells, (Hanahan, 2022; Hanahan & Weinberg, 2000, 2011) including uncontrolled proliferation, unresponsiveness to cell death induction and an altered metabolism. Cancer cells undergo a metabolic reprogramming favoring glycolysis and anaerobic respiration instead of oxidative phosphorylation, which is commonly known as “Warburg effect” (Locasale & Cantley, 2011; Warburg et al., 1927). Hexokinase (HK) catalyzes the first and irreversible step of glycolysis and thereby limits overall glycolytic activity. In mammals, the HK family consists of five members: HK1-4 and HKDC1 (HK domain containing 1). HKDC1 has an exceptionally low glucose affinity and, therefore, low hexokinase-activity under physiological conditions (Hayes et al., 2013). Thus, the exact function of HKDC1 mainly remains elusive. An increased *HKDC1* gene expression has been associated with chronic inflammation (Häsler et al., 2012; Quraishi et al., 2020; Weiser et al., 2018), metabolic diseases (Zapater et al., 2022) and various cancer types such as gastric cancer (Hu et al., 2023a; M. Wang et al., 2023; Yu et al., 2023; Zhao et al., 2023), liver cancer (Khan et al., 2022; Pusec et al., 2019; Y. Zhang et al., 2024), intrahepatic cholangiocarcinoma (Dong et al., 2022), endometrial cancer (Guo et al., 2022), breast cancer (X. Chen et al., 2019) or lung adenocarcinoma (X. Wang et al., 2020). In CRC, a splicing event in *HKDC1* was recently found to have significant associations with patient overall survival (Lian et al., 2020). Here, we aimed to investigate the function of HKDC1 for CRC progression and pathology by testing a linkage between HKDC1 expression and patient survival in CRC patients, generating and functionally profiling HKDC1-deficient human and murine cancer cell lines as well as normal and tumor-derived intestinal organoids *in vitro*, and finally analyzing the role of epithelial HKDC1 for intestinal carcinogenesis *in vivo* in mice. We show that deletion of HKDC1 in epithelial cells reduced cellular proliferation, restored sensitivity to mitochondria-mediated cell death, impaired mitochondrial respiration *in vitro* and, most important, ameliorated carcinogenesis in two independent mouse models - *Apc*^Min/+^ sporadic intestinal carcinogenesis and Caco-2 xenografts. Our findings therefore highlights a functional role of HKDC1 in CRC.

## Results

### Elevated *HKDC1* expression associates with prognostic significance in colon cancer

A role of HKDC1 had been shown for several cancer entities but its function within colon cancer remained unknown. We initially queried publicly available transcriptomic data from the human protein atlas (HPA, https://www.proteinatlas.org). Across different organs, *HKDC1* showed by far the highest expression within the gastrointestinal tract (Figure 1A) suggesting a potential role in intestinal function. Comparing various cancer pathologies using publicly available transcriptomic data from The Cancer Genome Atlas (TCGA, https://www.cancer.gov/ccg/research/genome-sequencing/tcga) project (The Cancer Genome Atlas Research Network et al., 2013), *HKDC1* expression was highest for colorectal cancer (average 19.3 FKPM) followed by pancreatic (average 10.7 FKPM), renal (average 8.1 FKPM), stomach (average 7.9 FKPM) and liver (average 6.2 FKPM) cancer (Figure 1B). Next, we compared the *HKDC1* expression in the intestinal mucosa between healthy controls and colon cancer (CRC) patients, which revealed that *HKDC1* expression was greatly increased in the transformed mucosa (Figure 1C), thus indicating its potential involvement in cancer development. Finally, using data from paired samples of normal mucosa and tumor tissue of the same CRC patients, we could further demonstrate an increased *HKDC1* expression in tumor compared to normal tissue, which indicates that *HKDC1* overexpression does not seem to depend on disease background but to be specific to the tumor tissue (Figure 1D).

**Figure 1.**
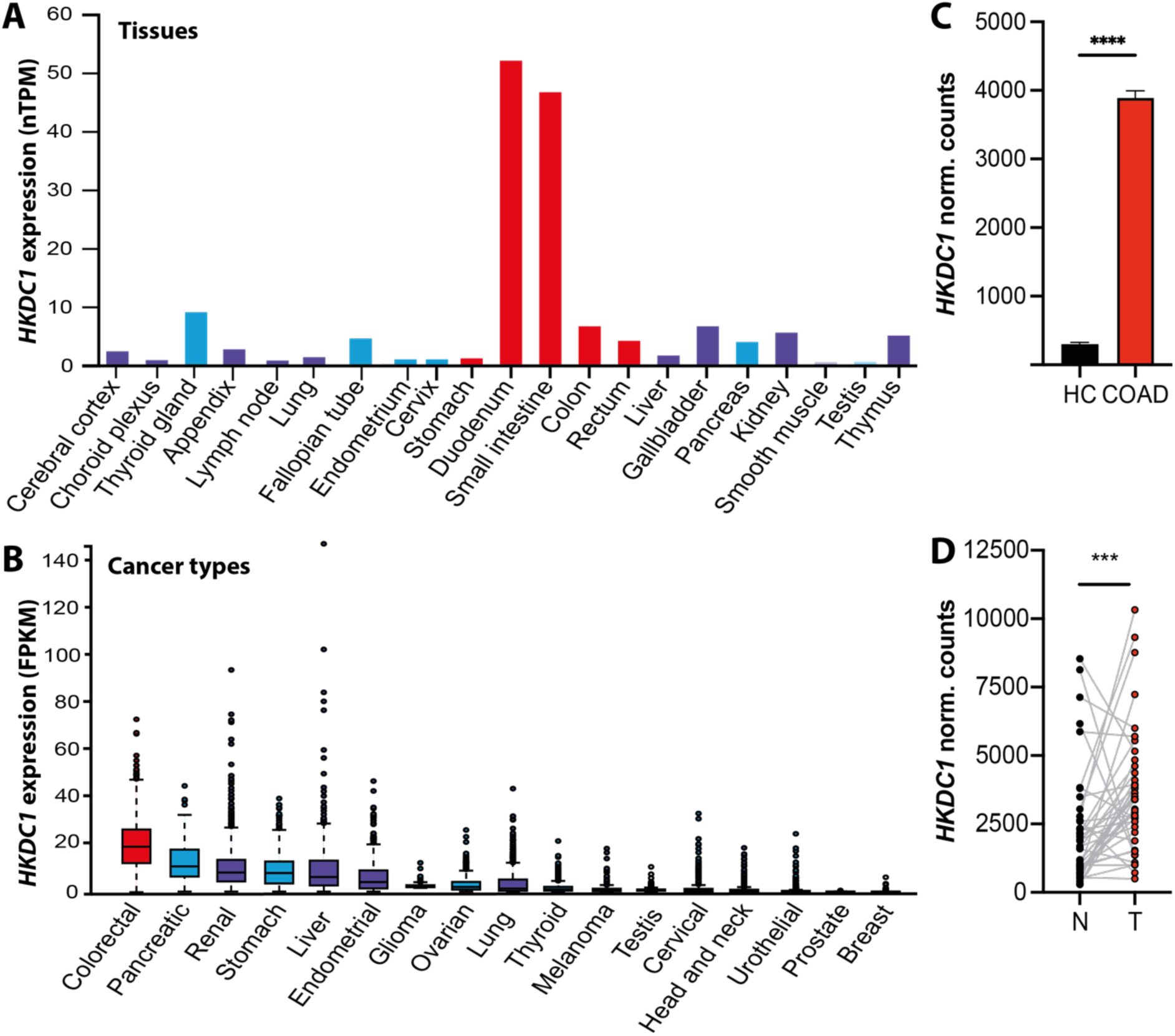
*HKDC1* expression is associated with colorectal cancer. **(A)** *HKDC1* expression as nTPM (normalized protein-coding transcripts per million) across different organs based on publicly available RNA-seq data from Human Protein Atlas (HPA). **(B)** *HKDC1* expression as FPKM (Fragments Per Kilobase per Million reads) from RNA-seq data of 17 cancer types of The Cancer Genome Atlas (TCGA, https://www.cancer.gov/ccg/research/genome-sequencing/tcga). The box plot depicts median values with the 0.25 and 0.75 percentiles. Outliers are identified if their values exceed 1.5 times the interquartile range. **(C)** *HKDC1* expression (normalized counts) in intestinal mucosa of healthy controls (HC) and CRC patients retrieved from publicly available databases The Genotype-Tissue Expression (GTEx, https://gtexportal.org, n = 454 samples) and TCGA (n = 651 samples). **(D)** *HKDC1* expression in paired normal (N) and tumor (T) tissue of n = 41 CRC patients from TCGA. ***: p < 0.001 as per Wilcoxon rank test. Data credits for **A** and **B**: Human Protein Atlas: www.proteinatlas.org, (Uhlén et al., 2015). The original plots are accessible at v23.proteinatlas.org/ENSG00000156510-HKDC1/tissue#rna_expression (A) and v23.protein atlas.org/ENSG00000156510-HKDC1/pathology (B).

### Loss of HKDC1 in colonic epithelial cells affects proliferation and cell death dynamics

Next, we aimed to investigate the function of HKDC1 for proliferation and cell death susceptibility using *in vitro* models, since abnormal or uncontrolled cell division and resistance to cell death represent fundamental hallmarks of cancer cells. To investigate these characteristics in a controlled *in vitro* environment we generated a murine (CMT-93) and a human (Caco-2) knockout colon epithelial cell lines using CRISPR technology. Both cell lines express *HKDC1* and the other HK family members *HK1* as well as *HK2*, but with differential patterns (Figure S1). Wildtype (WT) and HKDC1-deficient Caco-2 and CMT-93 cells were seeded with equal cell numbers and after 4 days cell growth was determined using multiple approaches. We either counted the number of generated cells within a cell forming assay or quantified protein content as a molecular measure of the cell number. HKDC1-deficient Caco-2 and CMT-93 cells had significantly reduced cell counts and protein content compared to WT cells (Figure 2A) indicating that HKDC1 contributes to cellular proliferation under homeostatic conditions. To investigate the function of HKDC1 in sensitivity towards cell death induction, WT and HKDC1-deficient Caco-2 and CMT-93 cells were stimulated with various agents that induce cell death, namely staurosporine, tumor necrosis factor (TNF), or interferon beta (IFN-β), and then analyzed for cell viability by fluorescence activated cell sorting (FACS). With the exception of staurosporine-treated Caco-2 cells, in general under all conditions, HKDC1-deficient Caco-2 and CMT-93 cells had a greater percentage of dead cells compared to WT controls (Figure 2B). Thus, loss of HKDC1 altered the cell death response *in vitro* in cancerous epithelial cells and especially affected mitochondria-dependent cell death.

**Figure 2:**
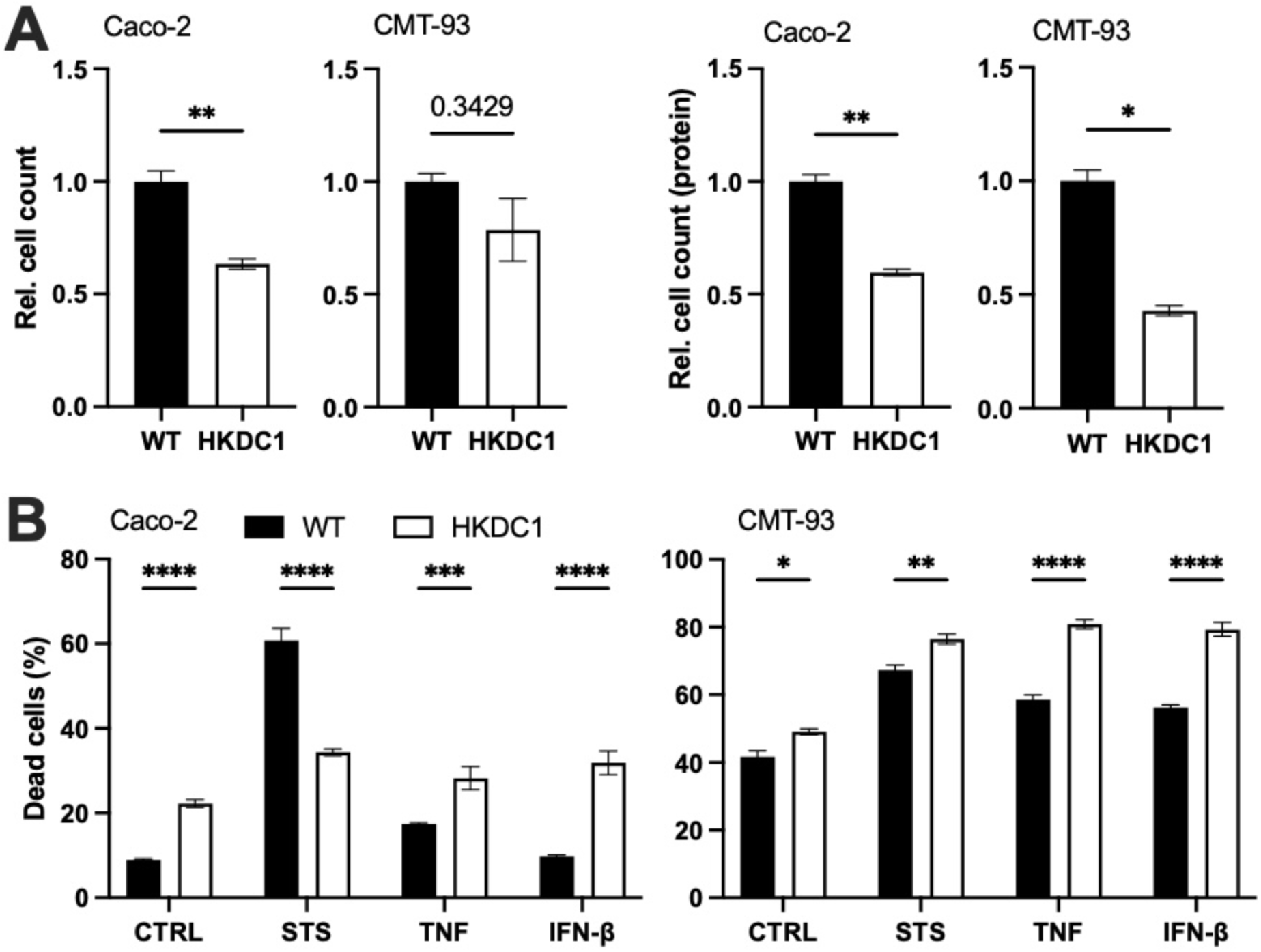
HKDC1-deletion alters proliferation and cell death. **(A)** Cell proliferation measured by two independent methods either as cell count or protein amount was reduced in HKDC1-deficient human Caco-2 and murine CMT-93 cells. n = 4 - 6 per group. *: p < 0.05, **: p < 0.05 as per Mann-Whitney-U-test. **(B)** WT and HKDC1-deficient Caco-2 and CMT-93 cells were stimulated with either staurosporine (STS, 10 µM for Caco-2 and 2 µM for CMT-93 cells), tumor necrosis factor (TNF, 500 ng/µl) or interferon beta (IFN-β, 1000 U/µl) for 24 hours and cell death was determined through zombie staining and FACS analysis. n = 5 per group. ***: p < 0.001 as per one-way ANOVA. All data is shown as mean ± SEM.

Next, we aimed to validate the *in vitro* findings made in cancerous epithelial cells (Caco-2 and CMT-93) *ex vivo* using intestinal epithelial organoids derived from *Hkdc1*^ΔIEC^ mice (Figure S2), a newly generated mouse line with a conditional knockout for *Hkdc1* specifically in intestinal epithelial cells (IECs) using the Villin::Cre/loxP system (Madison et al., 2002). Growth of these intestinal organoids was assessed using an organoid forming assay. Supporting the data from Caco-2 and CMT-93 cells, HKDC1-deficient organoids grew much slower and had a reduced overall cell mass compared to WT organoids (Figure 3A). The *Hkdc1*^ΔIEC^ mice were subsequently crossbred with tumor-bearing *Apc*^Min/+^ mice, a standard model for sporadic intestinal carcinogenesis (Uronis & Threadgill, 2009), to generate *Apc*^Min/+^-*Hkdc1*^ΔIEC^ mice. Notably, organoids derived from mice of the *Apc*^Min/+^ background grew much faster and as spherical cysts (Langlands et al., 2018) irrespective of the HKDC1 genotype. HKDC1-deficient *Apc*^Min/+^ organoids grew slower than WT *Apc*^Min/+^ organoids and also had a reduced overall cell mass (Figure 3B). In addition to proliferation of the intestinal organoids, we also assessed their sensitivity to cell death induction by challenging them with the same cell death inductors as the Caco-2 and CMT-93 cells earlier. In both the non-transformed and *Apc*^Min/+^ organoids, stimulation with the proinflammatory chemokines TNF and IFN-β did not induce cell death independent of the HKDC1 genotype (Figure 3C). The percentage of dead cells was reduced in untreated normal HKDC1-deficient compared to WT organoids but not in *Apc*^Min/+^ organoids. Notably, treatment with staurosporine, a known inducer of cell death through release of cytochrome c from the mitochondria into the cytosol (Ahlemeyer et al., 2002), induced cell death both in WT and HKDC1-deficient non-transformed organoids and also HKDC1-deficient *Apc*^Min/+^ organoids, whereas WT *Apc*^Min/+^ organoids were completely protected from staurosporine-induced cell death (Figure 3C). The transcriptional profiles from unstimulated WT and HKDC1-deficient *Apc*^Min/+^ organoids were surveyed by RNA sequencing. Gene ontology enrichment analysis revealed that proliferation and cell death processes were altered in HKDC1-deficient *Apc*^Min/+^ organoids compared to WT controls along with metabolic and immune processes (Figure 3D).

**Figure 3:**
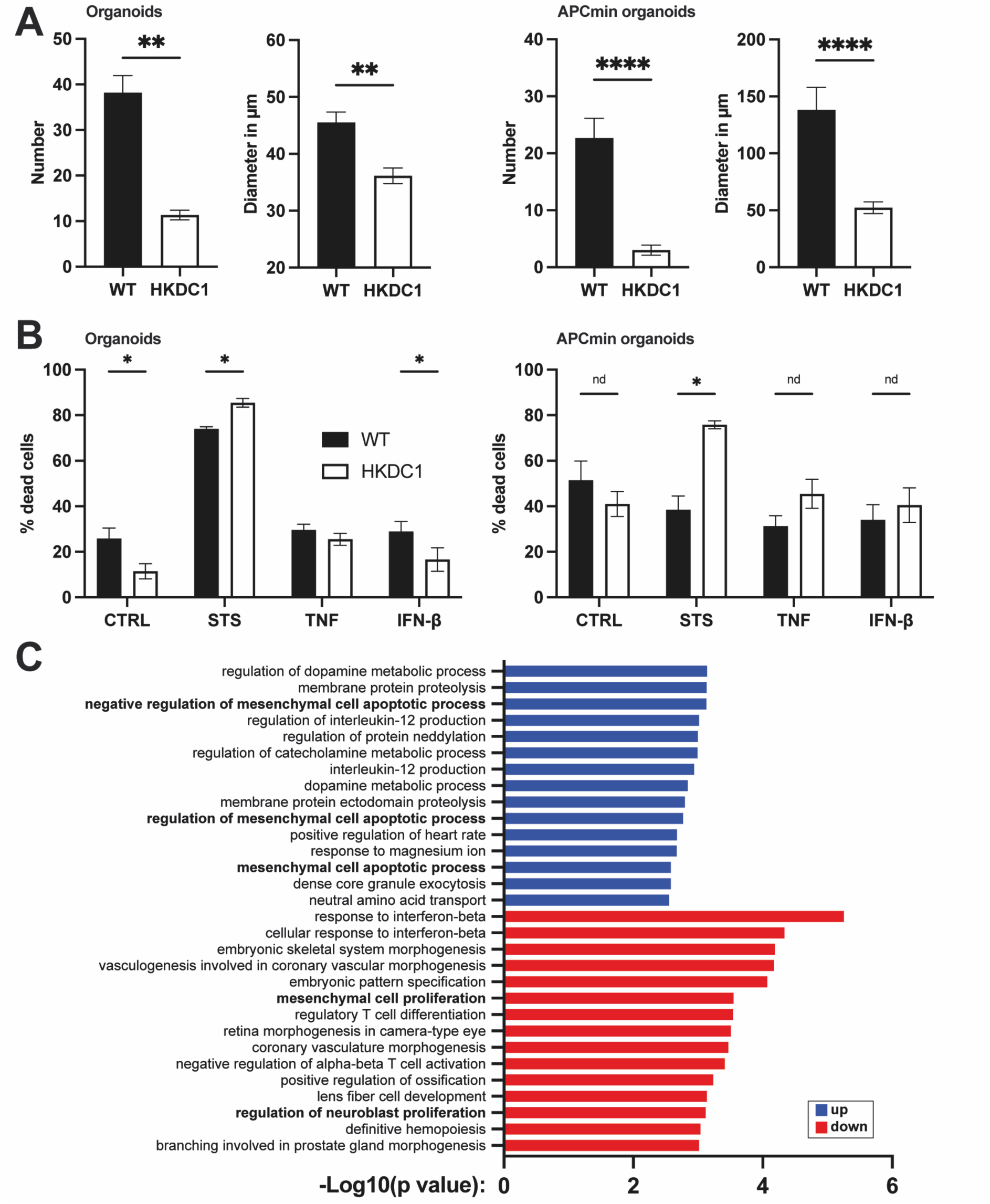
**Lack of HKDC1 reduced proliferation and altered cell death in murine organoids. (**A) HKDC1-deficient normal and tumorigenic *Apc*^Min/+^ organoids show a growth deficit as determined by organoid forming assay. Number and diameter of organoids five days after seeding (n = 6 - 12 per genotype and group). **: p < 0.01 and ****: p < 0.0001 as per Mann- Whitney-U-test. **(B)** Normal and tumorigenic *Apc*^Min/+^ organoids were stimulated with either staurosporine (STS, 2 µM), TNF (100 ng/ml), or IFN-β (50 ng/ml) for 24 hours and cell death was determined through zombie staining and FACS analysis. n = 5 per group. *: p < 0.05 as per Mann-Whitney-U-test. All data in **(A)** and **(B)** is shown as mean ± SEM. **(C)** Gene ontology (GO) biological process terms enriched among up- or downregulated genes in transcriptomes of organoids derived from *Apc*^Min/+^-*Hkdc1*^ΔIEC^ and littermate *Apc*^Min/+^-WT control mice. Top15 most enriched terms are shown. Proliferation and apoptosis functions are highlighted in bold.

### HKDC1 interacts with the mitochondrion and alters mitochondrial membrane potential

Seeking a more comprehensive understanding of the role of HKDC1 for cellular function, an HKDC1-targeted immunoprecipitation assay for possible interaction partners of HKDC1 was performed. In brief, protein lysates of scraped intestinal mucosa from H*kdc1*^ΔIEC^ and WT littermate mice were generated, incubated with an antibody against HKDC1 bound to dynabeads, washed, purified, and analyzed using LC-MS measurement. A *Hkdc1*^ΔIEC^ sample was included to increase specificity by only keeping candidate HKDC1 interaction partners that were found exclusively in WT samples. The identified proteins were linked with the LFQ (label free quantification) dataset (Van Puyvelde et al., 2022) and filtered to remove cytoskeletal proteins, which yielded a list of 34 candidate HKDC1 interaction partners (Table S1). A protein interaction network analysis revealed a distinct cluster of proteins associated with the regulation of mitochondrial membrane potential and pore activity (Figure 4A). This cluster included the other HK-isoforms HK1 and HK2 but also VDAC2 (Voltage-dependent anion channel 2) and VDAC3, SLC25A5 (Solute Carrier Family 25 Member 5 also referred to as ADP/ATP Translocase 2), ATP5A1 (ATP Synthase Subunit Alpha 1) and GFPT1 (Glutamine-Fructose-6-Phosphate Transaminase 1), which all function in mitochondrial metabolism.

**Figure 4:**
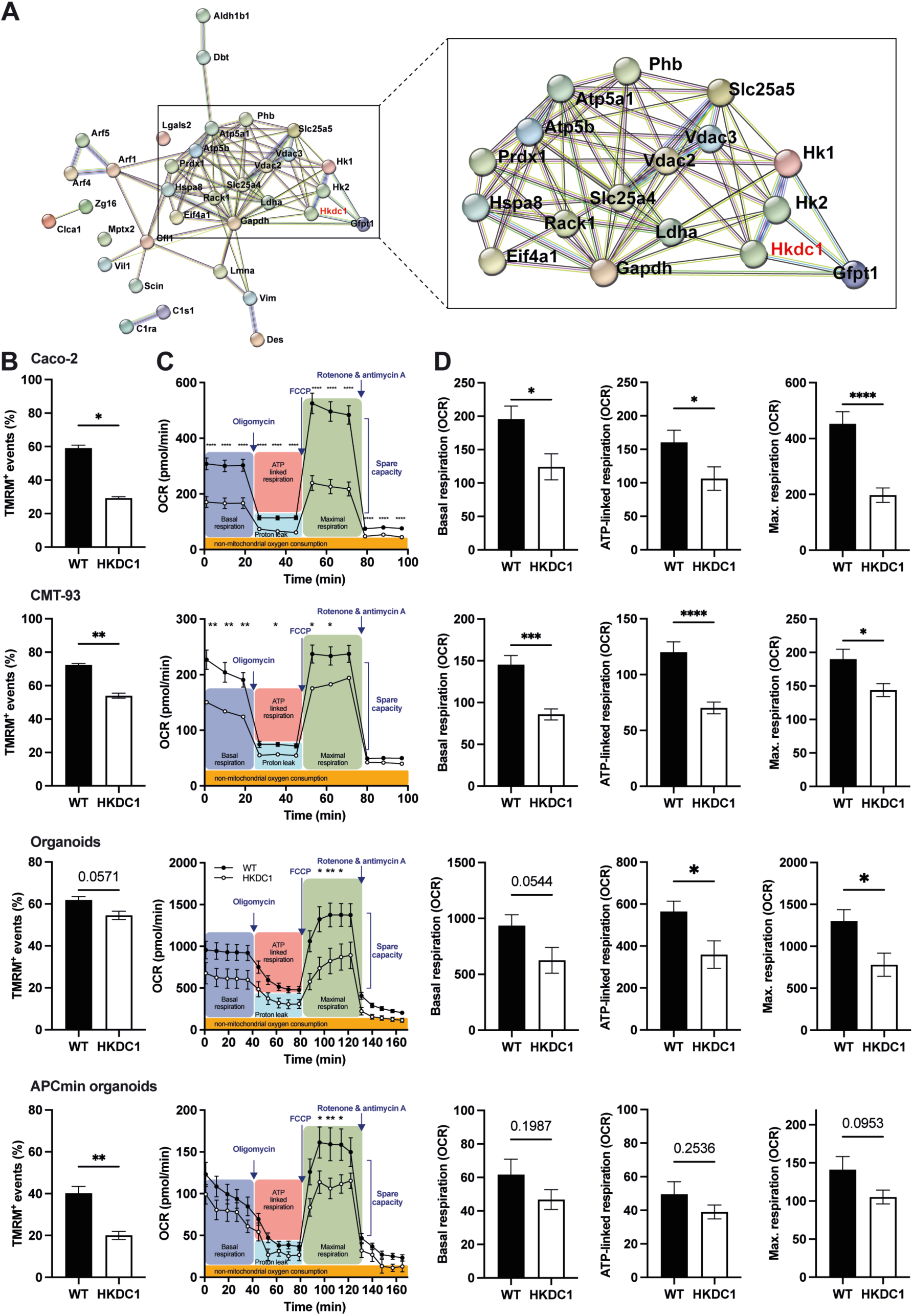
Lack of HKDC1 impairs mitochondrial function. **(A)** Network of HKDC1 protein interaction partners based on STRING analysis of identified proteins from HKDC1- immunoprecipitation of small intestinal mucosa highlighting a cluster of mitochondria-related proteins. Nodes represent individual proteins. Lines indicate known (cyan = curated databases, magenta = experimentally determined) and predicted (green = gene neighborhood, red = gene fusions, blue = gene co-occurrence) interactions. **(B-D)** Mitochondrial phenotyping in WT and HKDC1-deficient human Caco-2 and murine CMT-93 cells as well as intestinal normal and tumorigenic *Apc*^Min/+^ organoids. Deletion of HKDC1 reduced **(B)** mitochondrial membrane potential as per TMRM staining and FACS analysis and **(C)** oxygen consumption rate (OCR) as a measure of mitochondrial activity as determined by Seahorse Mito Stress metabolic analysis. **(D)** Basal as well as maximal respiration and ATP production as calculated from the depicted kinetic measurement. n = 9 - 10 per genotype and cell line/organoid. *: p < 0.05, **: p < 0.01, ***: p < 0.001 and ****: p < 0.0001 as per Mann- Whitney-U-test or two-way ANOVA. All data is shown as mean with SEM.

To test whether HKDC1 plays a role for mitochondrial function, we quantified the mitochondrial membrane potential in WT and HKDC1-deficient Caco-2 and CMT-93 cells as well as normal and *Apc*^Min/+^ organoids using staining with the MitoProbe tetramethylrhodamin-methylester (TMRM) and FACS analyses. TMRM signals were consistently reduced in HKDC1-deficient compared to WT cells for all models: Caco-2 (0.50- fold, p = 0.0159) and CMT-93 (0.75-fold, p = 0.0079) cells as well as normal (0.88-fold, p = 0.0571) and *Apc*^Min/+^ organoids (0.50-fold, p = 0.0079), indicating a reduced mitochondrial membrane potential in the absence of HKDC1 (Figure 4B). We then further profiled the role of HKDC1 for mitochondrial function by Seahorse extracellular flux XF analysis, which recently became available for organoids as well (Ludikhuize et al., 2021). Consistent with the findings from TMRM, HKDC1-deficient cells & organoids showed significantly lowered mitochondrial respiration (Figure 4C,D). Basal respiration along with ATP-linked and maximal respiration all were reduced in HKDC1-deficient cells highlighting a dysfunctional mitochondrial electron transport chain. Together, these data suggest a crucial role of HKDC1 in mitochondrial function.

### Epithelial deletion of HKDC1 ameliorated tumorigenesis

As HKDC1-deficient cancer cell lines and tumor organoids displayed a reduced proliferation, increased sensitivity to cell death induction and an impaired mitochondrial function, we next sought to test the role of HKDC1 in intestinal carcinogenesis *in vivo* using tumor-bearing *Apc*^Min/+^-*Hkdc1*^ΔIEC^ mice. Overall, body and organ weights (caecum, epididymal fat, spleen and liver) or measures (small intestinal and colon length) did not differ between *Apc*^Min/+^-*Hkdc1*^ΔIEC^ mice and WT littermate controls (Figure S3). Tumor counts in the small intestine and colon trended towards reduced counts in *Apc*^Min/+^-*Hkdc1*^ΔIEC^ mice compared to WT littermate controls, yet not reaching statistical significance (p = 0.1024 and p = 0.1973, respectively; Figure 5), which might be due to technical inaccuracies during the challenging identification of macroscopic tumors in the intestinal tissue. Supporting this data, histological analyses of colon sections revealed reduced numbers of Ki67-positive proliferating IECs (0.87-fold, p = 0.0377) along with slightly fewer apoptotic TUNEL-positive IECs (0.71-fold, p = 0.056) compared to littermate WT controls (Figure 5B-C). These data therefore support the former *in vitro* findings of reduced proliferation and an altered cell death sensitivity in cancerous colonic epithelial cells, namely Caco-2 and CMT-93 cell lines and *Apc*^Min/+^ tumor organoids, thus implying a role of HKDC1 in colon cancer. To further corroborate these data, we additionally performed a xenograft transplantation mouse model. To that end, WT and HKDC1-deficient Caco-2 cells were subcutaneously injected into the flank of immunocompromised NSG mice and tumor growth was monitored over time. 60 days post injection (dpi), mice transplanted with WT cells started to develop visible tumors, which continuously increased in volume until 70 dpi. In stark contrast, none of the mice transplanted with HKDC1-deficient Caco-2 cells developed any detectable tumor (Figure 5D-E). Thus, deletion of HKDC1 in Caco-2 cells completely abolished their capacity for tumor formation leading to full protection from cancer development. Taken together our data therefore highlights an important role of HKDC1 in intestinal cancer development.

**Figure 5:**
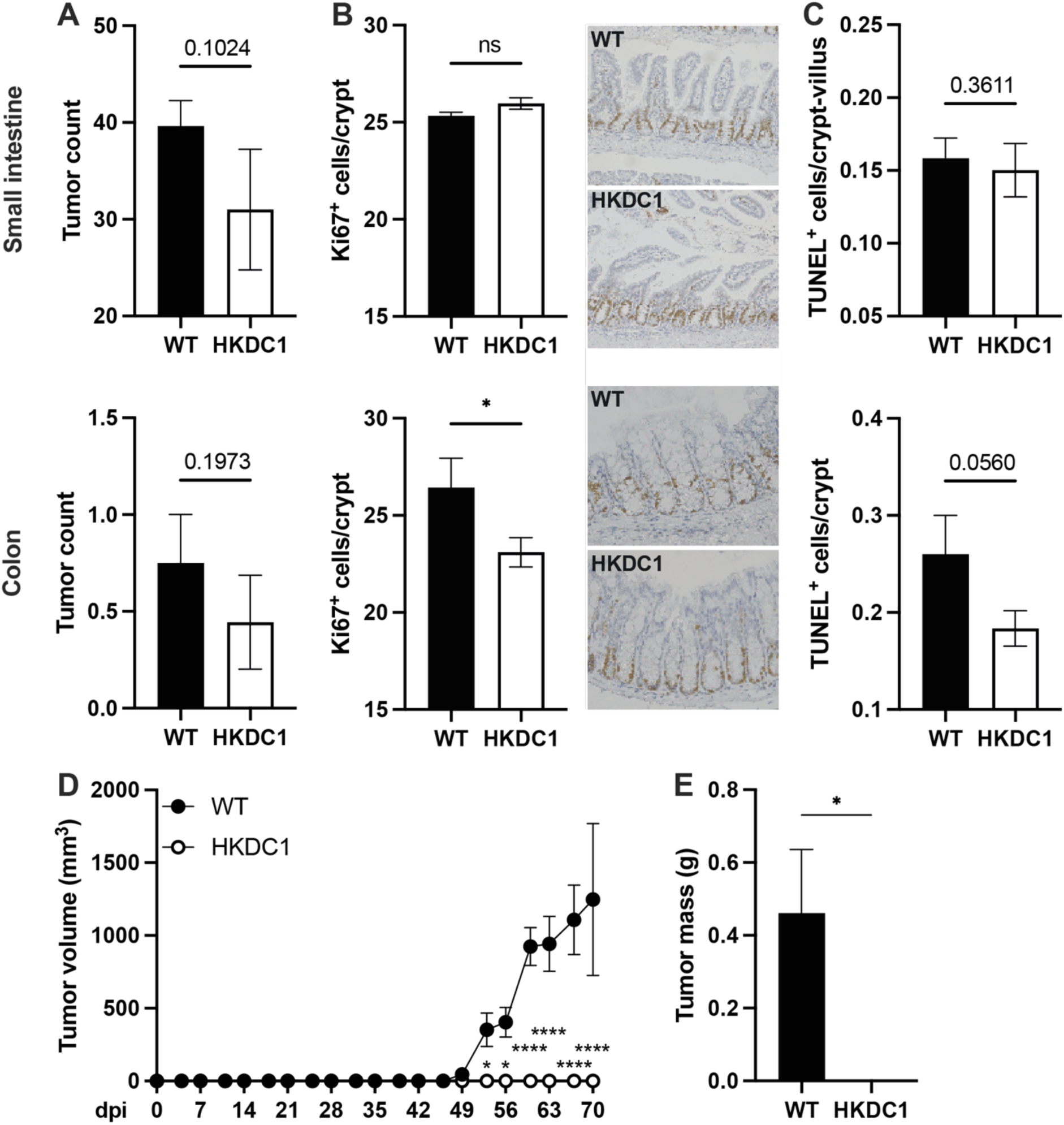
Epithelial deletion of *HKDC1* ameliorates intestinal carcinogenesis in two murine cancer models. (A-C) Sporadic intestinal tumorigenesis using the *Apc*^Min/+^ mouse model. 20- week-old tumor-bearing *Apc*^Min/+^-*Hkdc1*^ΔIEC^ mice were compared to WT littermate controls and analyzed for **(A)** tumor numbers, **(B)** proliferating (Ki-67 positive) and **(C)** apoptotic (TUNEL positive) IECs the small intestine and colon. n = 9 - 10 per genotype. *: p < 0.05 as per Mann-Whitney-U-test. **(D-E)** Xenograft model. NSG mice were injected with WT and HKDC1- deficient human Caco-2 and tumor volume (mm^3^) was monitored over time **(D)**. At day 70 post injection the experiment had to be stopped for ethical reasons, mice were killed and total tumor mass (g) was measured **(E)**. n = 4 mice per injection group. *: p < 0.05 and ****: p < 0.0001 as per two-way ANOVA. All data is shown as mean ± SEM.

## Discussion

Inhibition or deletion of HKDC1 protects from liver cancer (Khan et al., 2022; Y. Zhang et al., 2024) as well as gastric cancer (Yu et al., 2023), and dysregulation of HKDC1 had been associated to various cancer types (Q. Chen et al., 2020; X. Chen et al., 2019; J. Li et al., 2017; Lian et al., 2020; X. Wang et al., 2020), but its functional role for CRC remained unknown. In this study, we therefore aimed to clarify the function of HKDC1 for CRC. Our data revealed a linkage between *HKDC1* overexpression in tumor tissue compared to healthy tissue highlighting the importance of HKDC1 for cancer establishment and progression. We generated and functionally profiled HKDC1-deficient human and murine cancer cell lines as well as normal and tumor-derived intestinal organoids *in vitro*, and finally evaluated the role of epithelial HKDC1 for intestinal carcinogenesis *in vivo* in mice. These analyses showed that epithelial deletion of HKDC1 reduced cellular proliferation, restored sensitivity to mitochondria-mediated cell death and impaired mitochondrial respiration *in vitro* and mildly ameliorated colonic carcinogenesis in mice. Finally, deletion of HKDC1 in xenograft transplanted mice protected them from tumor formation. Our findings therefore highlight a functional role of HKDC1 in CRC development.

### HKDC1 contributes to cell proliferation and mitochondria-related cell death

Uncontrolled proliferation and unresponsiveness to cell death induction are two classic hallmarks of cancer cells (Hanahan & Weinberg, 2000). HKDC1-deficient Caco-2 and CMT-93 cells displayed reduced cell proliferation and an altered sensitivity to cell death induction (Figure 2) demonstrating that HKDC1 is functionally involved in these cancer hallmarks. Both cell lines essentially showed identical responses regarding proliferation and cell death induction with one notable exception: treatment with staurosporine reduced cell death in HKDC1-deficient Caco-2 cells whereas the percentage of dead cells was increased in HKDC1- deficient CMT-93 cells. These differential responses to staurosporine could be related to the differential expression pattern of HK isoenzymes in these two cell lines (Figure S1). In Caco-2 cells *HKDC1* was by far the most expressed HK, whereas in CMT-93 cells *Hk1* and *Hk2* were much more expressed than *Hkdc1*, thus a deletion of HKDC1 might be more effective in Caco- 2 than in CMT-93 cells as in CMT-93 cells the other HKs HK1 and HK2 could potentially compensate and carry out similar functions as HKDC1. We used the human and murine cell lines Caco-2 (Jumarie & Malo, 1991) and CMT-93 (Franks & Hemmings, 1978), which were derived from CRC patients and display phenotypic features of cancer cells. Therefore, these cell lines could exhibit distinct behaviors compared to primary cells, which could be a limitation of their use. We therefore also employed intestinal organoids derived from *Hkdc1*^ΔIEC^ mice and tumors of *Apc*^Min/+^-*Hkdc1*^ΔIEC^ mice, which validated the findings of reduced proliferation and an altered sensitivity to cell death induction (Figure 3) made in Caco-2 and CMT-93 cells. The definitive mechanism by which HKDC1 controls cell proliferation remains elusive, and further studies are needed to uncover the underlying molecular details. The altered sensitivity to an induction of mitochondria-related cell death in HKDC1-deficient cells could be linked to the association of HKDC1 to mitochondria and specifically proteins of the mitochondrial permeability transition pore (MPTP).

### HKDC1 associates with proteins of the mitochondrial permeability transition pore (MPTP)

Using immunoprecipitation, a powerful tool to study protein-protein interactions (Lin & Lai, 2017), we identified 34 proteins that potentially interact with HKDC1 (Figure 4A) including a cluster of proteins that are located at the mitochondrial membrane and function in mitochondrial metabolism, thus establishing a connection between HKDC1 and mitochondria. Mitochondria are central for cellular homeostasis, not only by orchestrating cellular bioenergetics but also by facilitating a metabolic switch regulating immune cell activation (Mills et al., 2017). Notably, the other HK-isoforms HK1 and HK2 were among the mitochondria-related cluster of potential HKDC1 interaction partners. As HKDC1 does not display a HK activity under physiological conditions due to its extremely low glucose-affinity (Hayes et al., 2013), HKDC1 could function as a “decoy”-HK that blocks the highly processive other HK-isoforms HK1 and HK2 through their direct interaction or maybe by competing to the available binding spots at the mitochondrial membrane, for example VDAC proteins, which were also among the identified HKDC1 interaction partners and are known to serve as mitochondrial anchoring points for HK1 (Jackson et al., 2015), HK2 (Baik et al., 2023) and HKDC1 (Pusec et al., 2023). Other mitochondria-related potential HKDC1 interaction partners included SLC25A5, ATP5A1 and GSPT1. SLC25A5 (also known as ANT2, adenine nucleotide translocator 2) plays a key role in maintaining mitochondrial membrane potential and inhibiting apoptosis by catalyzing the exchange of mitochondrial ATP with cytosolic ADP, which is a vital feature for cancer cells (Chevrollier et al., 2011). SLC25A5/ANT2 is believed to be a component of the MPTP although the exact composition of the MPTP remains unclear (Baines et al., 2005; Baines & Gutiérrez-Aguilar, 2018). Notably, we oberserved a reduced mitochondrial membrane potential in HKDC1-deficient cells and organoids, which is supported by previous publications (Q. Chen et al., 2020; Cui et al., 2024; Khan et al., 2022; Pusec et al., 2019). The reduced mitochondrial membrane potential in HKDC1-deficient cells could therefore be functionally related to its association with SLC25A5/ANT2 and regulation of the MPTP. Bioinformatic analyses previously highlighted the prognostic value of SLC25A5 in colon cancer (Y.-J. Chen et al., 2022). ATP5A1 is a subunit of the mitochondrial ATP synthase and implicated in promoting tumor growth during CRC (G. Zhang et al., 2022). GFPT1 is the first and rate-limiting enzyme of the hexosamine pathway and upregulation of GFPT1 promotes the proliferation of cervical cancer cells (Gong et al., 2021; D. Li et al., 2022). All of these HKDC1-interacting proteins are important for mitochondrial function, in particular the MPTP and mitochondrial membrane potential, but also cancer development, which therefore highlights a potential regulatory role of HKDC1 and therapeutic value of targeting HKDC1 in CRC.

### HKDC1 - a molecular target for colon cancer?

Experiments in cancer cell lines and intestinal organoids demonstrated reduced proliferation and an altered sensitivity to cell death induction in HKDC1-deficient compared to WT cells (Figure 2 and 3). An analysis of publicly available transcriptomic data of CRC patients revealed an overexpression of HKDC1 in tumor tissues (Figure 1). In combination with the reduced mitochondrial function in the absence of HKDC1 and the revealed interaction of HKDC1 with cancer-relevant & mitochondria-related proteins, our data clearly argues for a functional role of HKDC1 in CRC. Surprisingly, *Apc*^Min/+^-*Hkdc1*^ΔIEC^ mice showed only mildly improved disease phenotypes mainly in the colon (Figure 5). This surprising outcome could be based on the use of a conditional knockout of HKDC1 only in IECs leaving other cells such as immune or mesentery cells with functional HKDC1 that could potentially compensate the epithelial HKDC1 deletion. Furthermore, along the same lines, other HK-family members might compensate the deletion of HKDC1, although we did not detect any genotype-dependent differences in mRNA levels of the other HK-family members (Figure S3H). In *Apc*^Min/+^ mice, tumors mainly develop in the small intestine distinguishing it from human CRC, which primarily affects the colon. However, tumors also develop in the colon and this is where we saw the main protective phenotype of *Apc*^Min/+^-*Hkdc1*^ΔIEC^ mice. This would be in line with the *in vitro* findings that were made using cells lines of colonic human and murine cancerous epithelial cells. Even though, this data only suggested a mild effect of the deletion of HKDC1 in IECs upon tumor development, it still indicates the involvement of HKDC1 in tumor development. These data are in agreement with recent findings by Liu et al. (2024) demonstrating that HKDC1 functions as a glucose sensor within the tumor microenvironment (Liu et al., 2024). Mechanistically, HKDC1 stability is altered in response to glucose availability via a specific glucose sensing domain. This mechanism allows HKDC1 to promote tumor growth by interacting with prohibitin 2 (PHB2) leading to increased expression of pro-oncogenic molecules. These insights provide a deeper understanding of the intricate role of HKDC1 in cancer metabolism (Liu et al., 2024). To corroborate these data, an independent colon carcinoma-derived cell xenograft transplantation model was employed with WT and HKDC1 deficient Caco-2 cells. Importantly, mice transplanted with HKDC1-deficient xenografts did not develop any visible tumors over the course of the 10 weeks, whereas all mice receiving WT Caco-2 cells developed large tumors leading to termination of the experiment for ethical reasons. Thus, loss of HKDC1 in tumor tissue cells protected from tumor development. Prospectively, other *in vivo* models such as xenotransplantations of patient-derived organoids with targeted deletions of HKDC1 into immunocompromised mice could aid in a better understanding of the role of HKDC1 for CRC. Related results for other cancer entities confirmed a role of HKDC1 in carcinogenesis *in vivo* (X. Chen et al., 2019; Dong et al., 2022; Guo et al., 2022; Hu et al., 2023b; Khan et al., 2022; M. Wang et al., 2023; X. Wang et al., 2020; Yu et al., 2023; Y. Zhang et al., 2024; Zhao et al., 2023) providing further support for an important functional role of HKDC1 in CRC. A recent study revealed that HKDC1 promotes tumor immune evasion during immunotherapy with atezolizumab (anti-PD-L1) in hepatocellular carcinoma patients (Y. Zhang et al., 2024) adding further evidence for the therapeutic prospect of a HKDC1-targeted therapy during cancer. However, a future challenge will be to develop cancer cell-specific HKDC1-directed treatments given the physiological high presence of HKDC1 in the intestine to not harm non-transformed cells having a high HKDC1 expression.

## Conclusions

Our findings clearly identified HKDC1 as functioning in CRC development and demonstrated that HKDC1 plays a functional role in the regulation of proliferation, sensitivity to cell death and in particular mitochondrial metabolism. Using HKDC1 immunoprecipitation indicated a potential molecular mechanism by identifying a network of cancer-relevant mitochondrial proteins through which HKDC1 could impact cellular physiology. However, further studies are required to indeed demonstrate these molecular interactions and the feasibility and efficacy of HKDC1-targeted interventions for CRC.

## Methods

### Analyses of *HKDC1* expression in human CRC patients

RNA-seq data from normal tissues and tumor samples of human cancer patients were retrieved from multiple public databases: the Human Protein Atlas (HPA) publicly available at www.proteinatlas.org (Uhlén et al., 2015), The Cancer Genome Atlas (TCGA) available at https://www.cancer.gov/ccg/research/genome-sequencing/tcga (Hutter & Zenklusen, 2018; The Cancer Genome Atlas Research Network et al., 2013), The Genotype-Tissue Expression (GTEx) available at https://gtexportal.org/home (Lonsdale et al., 2013). Tissue RNA expression (HPA) were reported as nTPM (normalized protein-coding transcripts per million) corresponding to mean values of the different individual samples from each tissue. RNA-seq data from 17 cancer types (HPA) were reported as Median Fragments Per Kilobase of exon per Million reads (FPKM). RNA-seq data from healthy controls and CRC patients (GTEx and TCGA) as well as from paired normal and tumor tissue CRC patients (TCGA) were reported as normalized counts.

### Mice

All animal experiments were approved by the local animal safety review board of the federal ministry of Schleswig Holstein and conducted according to national and international laws and policies (approval numbers V242-56302/2018(100-11/18) and IX552-65205/2024(24- 4/24)). Mice were provided with autoclaved water and food ad libitum and maintained in a 12-hour light-dark cycle under standardized conditions (21 °C ± 2 °C with 60 % ± 5 % humidity) in the Central Animal Facility (ZTH) of the University Hospital Schleswig Holstein (UKSH, Kiel, Germany). Mice were housed in groups of up to five littermates in individually-ventilated cages (Green Line, Techniplast) under specific pathogen-free conditions. Mice were killed by cervical dislocation and tissues were removed for histological and molecular analyses.

Conditional-ready *Hkdc1*^tm1a(KOMP)Wtsi^ mice mice were purchased from the KOMP repository (https://www.komp.org, clone ID 799725, C57BL/6N-A^tm1Brd^ background). The mice were generated using a conditional-ready exon trapping strategy. In brief, a neomycin selection cassette was inserted into the *Hkdc1* genomic locus between exons 2 and 5. The cassette contained an FRT site followed by lacZ sequence, a loxP-flanked neomycin resistance gene with an internal FRT site and a third loxP site downstream of the targeted exons 3-4 (Figure S3). These *tm1a* offspring were bred with a flp-deleter strain purchased from Jackson Laboratory (https://www.jax.org, Stock Number: 011065) to remove the neomycin selection cassette and generate the final conditional allele *tm1c* (Figure S3). To create the *Hkdc1*^ΔIEC^ mouse line (alternatively termed *Hkdc1*^fl/fl^-*Villin*::Cre^+^ or “HKDC1”), we then crossed the conditional *tm1c* carrying floxed *Hkdc1* alleles with mice expressing the CRE recombinase under the control of the IEC-specific *Villin* promoter purchased from Jackson Laboratory (https://www.jax.org, Stock Number: 021504). As controls we used littermate *Hkdc1*^fl/fl^ mice not carrying any CRE recombinase, referred to as WT mice. For the *in vivo* analysis of HKDC1 function in intestinal carcinogenesis, *Apc*^Min/+^ (Adenomatous-polyposis-coli multiple intestinal neoplasia) mice were purchased from Jackson Laboratory (https://www.jax.org, Stock Number: 002020) and crossed with *Hkdc1*^ΔIEC^ mice to generate tumor-bearing *Apc*^Min/+^- *Hkdc1*^ΔIEC^ mice. *Apc*^Min/+^ mice are a standard model for sporadic intestinal carcinogenesis (Uronis & Threadgill, 2009). *Apc*^Min/+^ mice carry a mutated *Apc* gene encoding a nonfunctional truncated APC protein (Moser et al., 1995), which normally functions as a critical tumor suppressor by promoting the degradation of β-Catenin within the Wnt signaling pathway (Schneikert & Behrens, 2007) and thereby inhibiting activation of tumorigenic target genes such as *cmyc*.

### Xenograft mouse model

To generate subcutaneous xenografts, WT or HKDC1-deficient Caco-2 cells (each 2×10^6^ cells in 100 µl PBS) were injected into the right flank of 8- to 12-week old male NOD.Cg-Prkdc^SCID^ Il2rg^tm1Wjl^/SzJ (NSG) mice purchased from Charles River Germany (Czulkies et al., 2017). Viability of the injected Caco-2 cells was determined before and after injection using plating and PI staining (Logos Biosystems, catalogue number F23001, Lot AP0BBH2901). Mice were checked for disease symptoms multiple times per week. Tumor growth and volume were traced by measuring a palpable material using a caliper. For ethical reasons, the experiment had to be discontinued at day 70 as tumors in the WT group reached the endpoint volume of 1500 mm^3^. Mice were killed by cervical dislocation and the subcutanous tumors were dissected, measured to determine tumor volume and weight to determine tumor mass.

### Histology and immunostaining

Intestines were flushed with PBS, cut open longitudinally and rolled from distal to proximal. Swiss rolls were fixed in in 10% formalin solution (ThermoFisher Scientific) over night at 4 °C and then embedded in paraffin. 5 µm thick sections were cut and stained with hematoxylin and eosin (H&E) or subjected to immunostaining using the Vectastain ABC kit (Vector Laboratories) including antigen retrieval in boiling citrate buffer. Primary antibodies were incubated overnight. For immunostaining of Ki67 we used a 1:500 diluted mouse anti-Ki67 antibody (BD Biosciences, cat.no. 556003). The TUNEL assay was performed using the ApopTag Plus Peroxidase In Situ Apoptosis Detection Kit (Merck Millipore) according to the manufacturer’s instructions. Slides were visualized using a Zeiss Imager Z1 microscope (Zeiss) and pictures were taken using ZEN pro software (Zeiss, version 3.4). Proliferative and apoptotic cells were quantified by counting Ki67- and TUNEL-positive cells in at least 30 randomly selected crypts of a sample in a blinded fashion by keeping the mouse identity masked during the counting process.

### Isolation and culture of intestinal organoids

Organoids were generated from intestinal crypts of *Hkdc1*^ΔIEC^ and *Apc*^Min/+^-*Hkdc1*^ΔIEC^ mice following established procedures as described before (Sato et al., 2011). The desired amount of cells was plated in a 50 µl drop of a 1:1 mixture of Advanced DMEM/F12 and Matrigel (BD). ENR-conditioned medium consisted of 70 % (*v*/*v*) 2 × basal medium (Advanced DMEM/F12 supplemented with HEPES [1M], Glutamax [100×], Penicillin/streptomycin 10,000 U/mL [1:50] and N-Acetylcysteine [500 mM]), 10 % (*v*/*v*) Noggin-conditioned medium and 20 % R- Spondin-conditioned medium. Noggin- and R-Spondin-conditioned media were prepared as described below. Organoids were cultivated in 24 well plates at 37°C with 5 % CO2 atmosphere in Matrigel (BD) with ENR-conditioned medium supplemented with 0.1 % human recombinant EGF (50 µg/mL). Medium was changed every two to three days and organoids were passaged every four to eight days. Organoids were cultured for at least two passages before being used for experiments.

### Organoid forming assay

Organoids were resuspended in 1 ml TrypLE Express (Invitrogen) supplemented with 10 µM Y-27632 (ThermoFisher Scientific) and incubated at 37 °C to create single cells. The desired cell number (5000 cells per 20 µl and well) was cultured in a 24 well plate. After a 5-day growth period, organoids were counted and diameters measured using ZEN software (version 3.4).

### Cell culture

Caco-2 and CMT-93 cells were purchased from DSMZ (ACC-169) and ATCC (CCL-223). Cells were cultured at 37 °C and 5 % CO2 in their respective media (Caco-2: MEM + 20 % Fetal bovine serum (FBS), CMT-93: DMEM + 10 % FBS) until a fully confluent cellular monolayer was established. Cells were stimulated at a confluency of 70 %.

HEK Noggin cells (Miyoshi & Stappenbeck, 2013) kindly provided by Prof. Zeissig (CRTD, Dresden University, Germany) were cultured for three cycles in DMEM containing 10 % FBS and 10 µg/ml Puromycin (Sigma-Aldrich). Afterwards, cells were diluted 1:20 in DMEM + 10 % FBS and cultured for four days. Then the supernatant was collected and centrifuged to eliminate cells. This was repeated after culturing for four more days and both batches combined. HEK R-Spondin-1-producing cells (Miyoshi & Stappenbeck, 2013) kindly provided by Prof. Zeissig (CRTD, Dresden University, Germany) were cultured for three cycles in DMEM containing 10 % FBS and 300 g/ml Zeocin (Thermo Fischer Scientific), following a 1:20 dilution in DMEM + 10 % FBS. The cells were cultured for four days, media was collected and centrifuged to remove cells. This was repeated after culturing for four more days and both batches combined.

### Generation and culture of HKDC1-deficient Caco-2 and CMT-93 cells

HKDC1-deficient cell lines were generated by CRISPR/Cas9 technique with plasmids that were assembled using the GeneArt™ CRISPR Nuclease Vector with CD4 Enrichment Kit (Thermo Fisher Scientific) according to the manufacturer’s instructions with the following primers: CRISPR_Caco-2_For TTC CCG CGG ATG ATT TCA TTG TTT T and CRISPR_Caco-2_Rev AAT GAA ATC ATC CGC GGG AAC GGT G, CRISPR_CMT-93_For CCT GTA TCA CAT GCG GCT CTG TTT T and CRISPR_CMT-93_Rev AGA GCC GCA TGT GAT ACA GGC GGT G. Correct plasmid sequences were validated using inhouse Sanger sequencing. Caco-2 or CMT-93 cells were transfected with the respective *HKDC1* or *Hkdc1* CRISPR plasmid using the Lipofectamine 3000 reagent kit (Thermo Fisher Scientific). Positive clones were purified using the Dynabeads® CD4 Positive Isolation Kit (Thermo Fisher Scientific) and thorough washing until cells directly bound to beads remained and no secondary bindings were present as monitored by stereomicroscopic evaluation. Beads were detached, cell numbers were counted and single cells were seeded per well of a 96-well plate. Single colonies were picked, cultured on a larger scale and screened for the absence of HKDC1 using western blots to generate a monoclonal population of HKDC1-deficient Caco-2 and CMT-93 cell clones. Caco-2 or CMT-93 cells that were also subjected to the CRISPR transfection and selection procedure, but which still showed a HKDC1 band as per western blot analysis were used as WT controls.

### *In vitro* cell formation assays

Proliferation of WT and HKDC1-deficient Caco-2 and CMT-93 cells was assesed using multiple methodologies: either by estimating cell density or using the protein content of the sample as a measure for the cell count. To estimate the cell density, Caco-2 or CMT-93 cells of the regular culture were detached and initially counted using Cellometers Auto T4 Plus (Heraeus). A fixed cell number (CMT-93: 1×10^6^, Caco-2: 1.5×10^6^) was seeded on 24-well plates and four days after seeding the cell count was determined again to monitor the increase in cell mass. Alternatively, protein was extracted as described below as a molecular measure of the cell count. Proteins were extracted from cell pellets or scrapings of small intestinal mucosa from *Hkdc1*^ΔIEC^ mice and WT littermate controls. Samples were lyzed in ice-cold RIPA buffer containing protease and phosphatase inhibitors and were homogenized by sonication (cells) or bead beating (tissue) in tissue lyser (Qiagen). After centrifugation, the supernatant was used for measuring protein concentration. The entire procedure was conducted at 4 °C or on ice. Protein was then quantified using the BCA assay (BioRad) according to the manufacturer’s instructions on a M200 Pro microplate reader (Tecan).

### Cell death quantification

Cell ceath was assesed using a viability staining based on zombie red (BioLegend), which selectively stains dead cells due to their compromised cell membranes, coupled to FACS analysis. Caco-2 and CMT-93 cells or organoids were cultured as described above in 24-well plates. Stimulations with staurosporine (10 µM for Caco-2, 2 µM for CMT-93 cells and for organoids), TNF (100 ng/ml) or IFN-b (50 ng/ml) for 24 hours were performed to induce cell death. Stimulated cells and organoids were washed and dissociated into single cells as described. The cells were then resuspended in 100 µl FACS wash buffer (1 % FCS in PBS), transferred to a 96-well plate and stained with zombie red (BioLegend, 1:1000 dilution in FACS wash buffer) for 30 min at room temperature in the dark. After washing, the cells were resuspended in 100 µl FACS wash buffer and subjected to FACS analysis using the Spectral Cell Analyzer (SONY). The percentage of dead cells was determined based on Zombie red intensity after excluding doublets and debris in the Sony SA3800 software (version 2.0.5).

### Mitochondrial membrane potential

Mitochondrial membrane potential was assessed using the MitoProbe tetramethylrhodamin- methylester (TMRM) Assay-Kit (Thermo Fisher Scientific). The positively charged TMRM translocates through the mitochondrial membrane accumulating in the negatively charged mitochondrial matrix (Creed & McKenzie, 2019) and a reduced TMRM staining therefore indicates a disrupted mitochondrial membrane potential. Caco-2 and CMT-93 cells were cultured overnight and organoids were harvested on the day of staining. TMRM was added a final concentration of 20 nM and incubated for 30 minutes at 37 °C, 5 % CO2. After a brief wash, Caco-2 and CMT-93 cells were trypsinated whereas organoids were incubated with TrypLE Express (Thermo Fisher Scientific) 15 minutes to create single cells. The single cell solutions were centrifuged at 300 x g for 5 minutes, washed and resuspended in FACS wash buffer. All cells were measured using the Spectral Cell Analyzer (SONY). The percentage of TMRM-positive cells was determined after excluding doublets and debris in the Sony SA3800 software (version 2.0.5).

### Seahorse metabolic assays

The mitochondrial activity of Caco-2 and CMT-93 cells as well as intestinal organoids were measured in real-time using the Seahorse XF Cell Mito Stress Test Kit according to the manufacturer’s instructions on a Seahorse XFe24 Analyzer (Agilent Technologies). Three independent experiments were performed with n=9 replicates. For Caco-2 or CMT-93, 4×10^4^ cells were seeded 24 hours for the assay. For intestinal organoids, 5000 cells were seeded as 20 µl drop in pure Matrigel onto the Agilent Seahorse XF24 cell culture microplate before 400 µl of organoid culture medium was added and the assay was performed as described before (Ludikhuize et al., 2021).

### RNA isolation and qPCR

Total RNA was extracted from cell pellets, organoids or intestinal tissue using the RNeasy Mini Kit (Qiagen) according to the manufacturer’s protocol. RNA concentration was measured using a NanoDrop ND-1000 spectrophotometer (PeqLab Biotechnologie). Total RNA was reverse transcribed to cDNA using the Maxima H Minus First Strand cDNA Synthesis Kit (ThermoFisher Scientific). qPCR was carried out using SYBR Select Master Mix (Applied Biosystems) according to the manufacturer’s instructions on a Viia 7 Real-Time PCR System (ThermoFisher Scientific). Expression levels were normalized to *Actb* (β-actin). Primer sequences for qPCR are listed in Table S2.

### Transcriptional profiling by RNA sequencing

Total RNA from small intestinal tissue, tumors and intestinal organoids of *Apc*^Min/+^-*Hkdc1*^ΔIEC^ and WT littermate mice was extracted using the RNeasy Mini Kit (Qiagen) according to the manufacturer’s protocol. RNA concentration and integrity were analysed using a TapeStation 4200 System (Agilent) and a Qubit 4 fluorometer (ThermoFisher Scientific). RNA-sequencing libraries were prepared using the TruSeq® RNA seq Library Prep Kit v2 according to the Illumina TruSeq® messenger (mRNA) sequencing protocol. The RNA-seq libraries were sequenced on an Illumina NovaSeq 6000 sequencer (Illumina, San Diego,CA) with an average of 15 million paired-end reads (2x 150 bp) at IKMB NGS core facilities. Adapter sequences were removed from the raw reads using cutadapt (v2.8) with a minimum overlap of 3 bp and a maximum error tolerance of 10% mismatches (TrueSeq forward adapter and TruSeq reverse complement universal adapter). Additional 3’- end quality trimming was performed to a minimum Phred score of 25, and poly-G ends were trimmed to address dark-cycle issues associated with Illumina’s two-color chemistry, utilizing the cutadapt option --nextseq-trim=25. An additional quality filtering step was applied using PrinSeq Lite (v0.20.4) to achieve a mean read quality of at least Phred score 30 across all bases, with a maximum of 5 unknown base calls and a minimum read length of 30 bp. Post-QC read qualities were visually inspected using FastQC (v0.11.7). The filtered reads were then mapped to the Mus musculus reference genome GRCm38, released by the European Bioinformatics Institute in version 99 (February 2020), using Hisat2 (v2.1.0) software. Only uniquely mapped reads were retained, utilizing SamTools (v1.9) with the -F 256 flag. Gene and exon abundances were then counted for properly mapped read pairs, with strandedness information (-s 2) taken into account, using the “featureCounts” tool from the subread software (v2.0.1). Samples that did not meet the QC standards were excluded from further analysis. Count data was analyzed in R (version 4.4.0 15/11/2024 17:22:00) using the DESEq2 R package (version 1.44.0 (Love et al., 2014)) to obtained differentially expressed (DE) genes. Genes with less than 10 counts across all samples were removed prior to the analysis. After comparison of KO vs WT samples within the respective tissues, genes with adjusted p-value (FDR) below 0.05 were retained for further analysis. Subsequently, a Gene Ontology Biological Process (BP) enrichment was performed in R with the ClusterProfiler package (version 4.12.0 (Wu et al., 2021)) using the annotations in the org.Mm.eg.db package (version 3.19.1 (Carlson, 2024)); genes used for the DE analysis were used as the gene universe. Genes with positive and negative log2 fold changes were enriched separately for each comparison.

### HKDC1 immunoprecipitation and liquid chromatography-mass spectrometry

Protein lysates from small intestinal mucosal scrapings of HKDC1^ΔIEC^ and WT littermate mice were used for Immunoprecipitation (IP). 1 µg of anti-HKDC1 antibody (Abcam, ab2279278) was bound to 50 µl Dynabeads (Thermo Fisher Scientific, 10004D) by incubation at 4 °C for 1 hour and then crosslinked in 250 µl 5 M bis(sulfosuccinimidyl)suberate for 30 min at room temperature. The crosslinking reaction was stopped by adding 12.5 µl quenching buffer (1M Tris-HCl pH 7.5) and incubation at room temperature for 15 minutes. After washing, 200 µl protein lysate was added to the crosslinked beads and incubated at 4 °C over night. The beads were washed and transferred into a fresh tube. Elution was performed by adding 22 µl elution buffer (50mM Glycine pH 2.8) and incubating for 5 minutes at room temperature. 10 µl of Tris pH 7.5 was added and samples were stored at -20 °C. The IP eluates were analyzed using liquid chromatography-mass spectrometry (LC-MS). Briefly, sample cleanup was performed according to the SP3 protocol (Hughes et al., 2019). The samples were first reduced with 10 mM dithiothreitol at 56 °C for 1 hour, followed by alkylation with 50 mM iodoacetamide in the dark at 20 °C for 30 minutes. Proteins were then precipitated onto hydrophilic and hydrophobic Sera-Mag SpeedBeads (carboxylate-modified magnetic beads, hydrophilic: 45152105050250, hydrophobic: 65152105050250, GE Life Sciences) using 6-fold volume of ethanol. After washing three times with 80 % ethanol, the samples were digested with trypsin (approximated enzyme:protein ratio of 1:40) overnight at 37 °C. Peptides were acidified to pH 2-3 using trifluroacetic acid (TFA) prior to LC-MS analysis on a Dionex U3000 UHPLC system (ThermoFisher Scientific) equipped with a column chromatography setup coupled to a Q Exactive Plus Orbitrap MS (ThermoFisher Scientific) using a nanospray ion source with 1.7 kV spray voltage and 250°C capillary temperature with a 20 µm Tip emitter (MS Wil). Peptides were concentrated and washed onto a trap column (75 μm × 2 cm, 2 μm C18 resin, 100 Å; Acclaim PepMap100, Thermo Fisher Scientific) for 5 min with 2% ACN and 0.05% aqueous TFA at a flow rate of 30 µl/min. Subsequently, peptides were separated on an analytical column (75 µm x 50 cm, 2 μm C18 resin, 100 Å; Acclaim PepMap100, Thermo Fisher Scientific) at 300 nl/min and separated using a 60 min linearly increasing gradient of LC solvent B (80 % acetonitril (ACN), 0.1 % formic acid (FA)) in LC solvent A (0.1 % FA). The linear gradient was followed by a sharp increase to 90 % solvent B for 5 min, an isocratic 10 min washing step and finally a column equilibration with 5 % B for 12 min. After a delay time of 5 min, the MS acquisition program consisted of a full-scan MS (range: 300 – 1,800 m/z, resolution: 60,000, automatic gain control (ACG) target: 3e6, maximal injection time (IT): 50 ms) with the top 10 MS/MS acquisition of the most intense ions using a 2.0 m/z isolation (resolution: 15,000, AGC target: 1e5, maximal IT: 50 ms). Ions of unassigned, +1, and >+8 charge states were excluded. For fragmentation, higher-energy collisional dissociation (HCD) was utilized with a normalized collision energy (NCE) of 27, Dynamic exclusion (20 s) and lock mass (445.12003 m/z) were enabled. Each sample was analysed using two technical replicates. Raw MS data were searched against the UniProt reference proteome of *Mus musculus* strain C57BL/6J (55.315 entries, 28. March.2022) and common contaminants (cRAP, contact/dust and laboratory contaminants; 42 entries) using Proteome Discoverer (version 2.2.0.388; ThermoFisher Scientific). Percolator was used for posterior error calculation (Käll et al., 2008) combining the results of the database searches restricted by q-value to FDR ≤ 0.01. A given protein was considered as “identified” when a valid MS^2^ spectrum was available for at least one of the peptides belonging to that protein. Detailed settings are listed in Table S3. Structural proteins such as cytoskeletal proteins (myosin, actin, tubulin, tropomodulin), keratins, histones and ribosomal proteins along with immunoglobulin proteins and proteins having a coverage of less than 25% were removed from further analyses. To generate a protein-protein interaction network and to visualize functional hubs among the identified HKDC1-interaction partners, the Search Tool for the Retrieval of Interacting Genes (STRING: https://string-db.org/ (version 12.0, accessed on 22 February 2024)) was used (Szklarczyk et al., 2023) in medium confidence (0.400) mode.

### Statistics

General statistical analyses were performed using the GraphPad Prism 9 (GraphPad Software Inc., La Jolla, USA). For pairwise comparisons the Mann-Whitney-U-test was used, whereas for multiple comparisons one-way or two-way ANOVA with false discovery rate (FDR) correction were performed. Data are shown as mean ± standard error of the mean (SEM). A p-value < 0.05 was considered as significant (*). A p-value < 0.01 was considered as strongly significant (**) and p-values of p < 0.001 (***) and p < 0.0001 (****) as highly significant.

---

**Figure S1:**
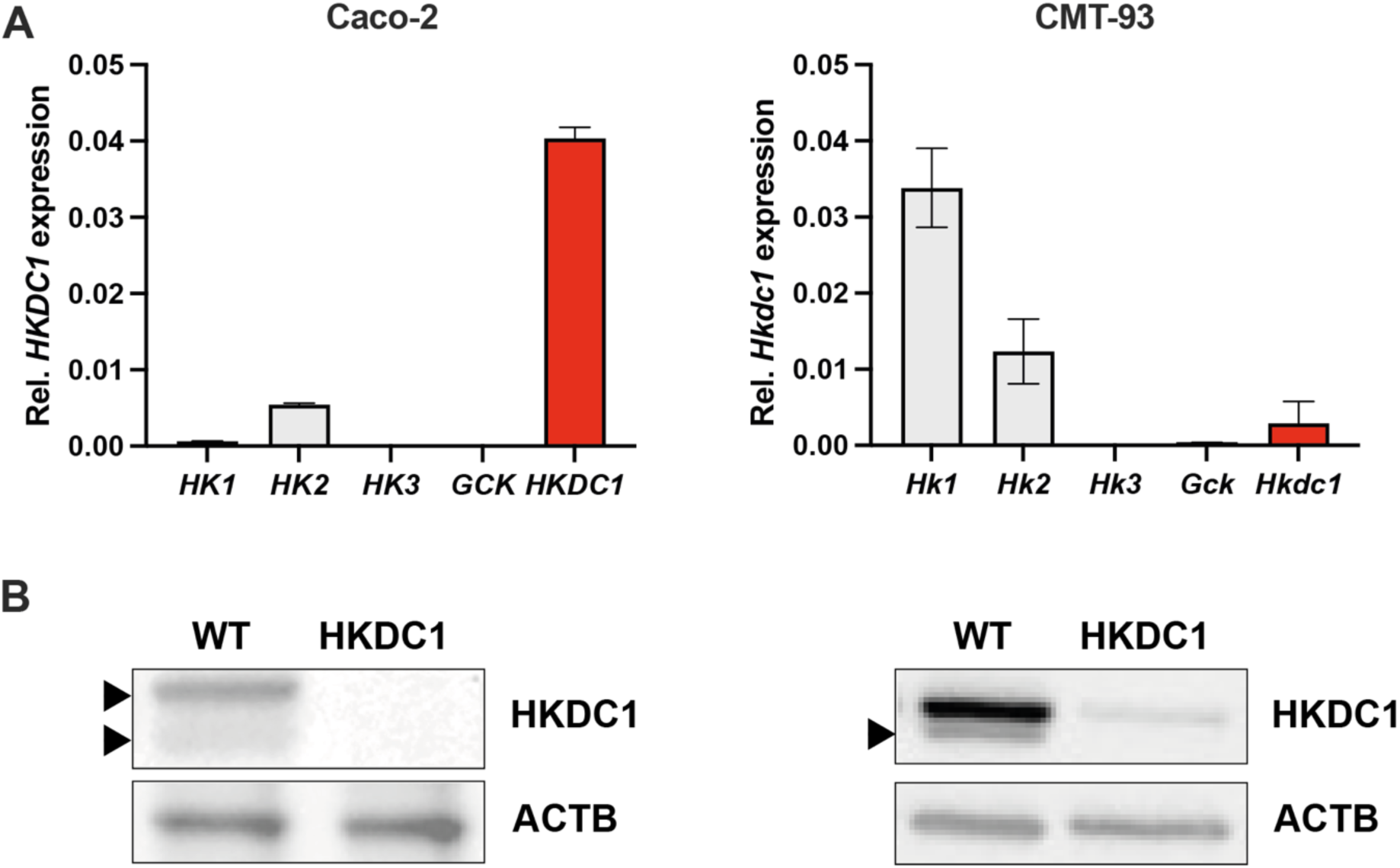
Expression of all hexokinases in the used cell lines. **(A)** Relative gene expression of all HK family members in human Caco-2 cells and murine CMT-93 cells normalized to beta Actin (*ACTB/Actb*) expression. GCK = Glucokinase. n = 5 per group. **(B)** Westernblot demonstrating functional HKDC1 deletion in human Caco-2 cells and murine CMT-93 cells.

**Figure S2:**
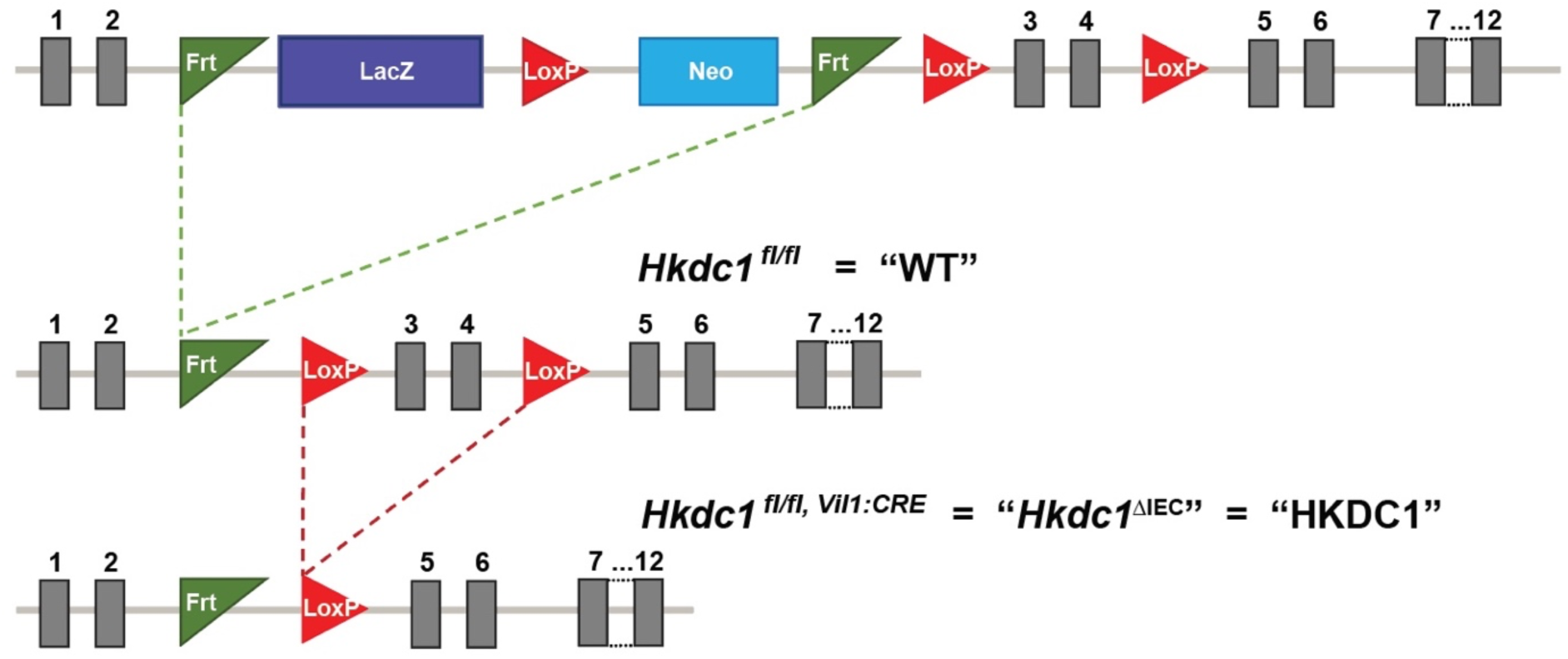
Strategy for generating *Hkdc1*^ΔIEC^ mice. IEC-specific HKDC1-deficient mice were generated by crossing conditional-ready *Hkdc1*^tm1a^ with Flp-deleter mice to create *Hkdc1* floxed (Hkdc1^fl/fl^) mice, which then were bred with Villin::CRE transgenic mice expressing CRE recombinase under the control of the IEC-specific *Villin1* promoter, resulting in *Hkdc1*^ΔIEC^ mice carrying a specific conditional deletion of *Hkdc1* exons 3-4 only in IECs and a non-functional HKDC1 protein.

**Figure S3:**
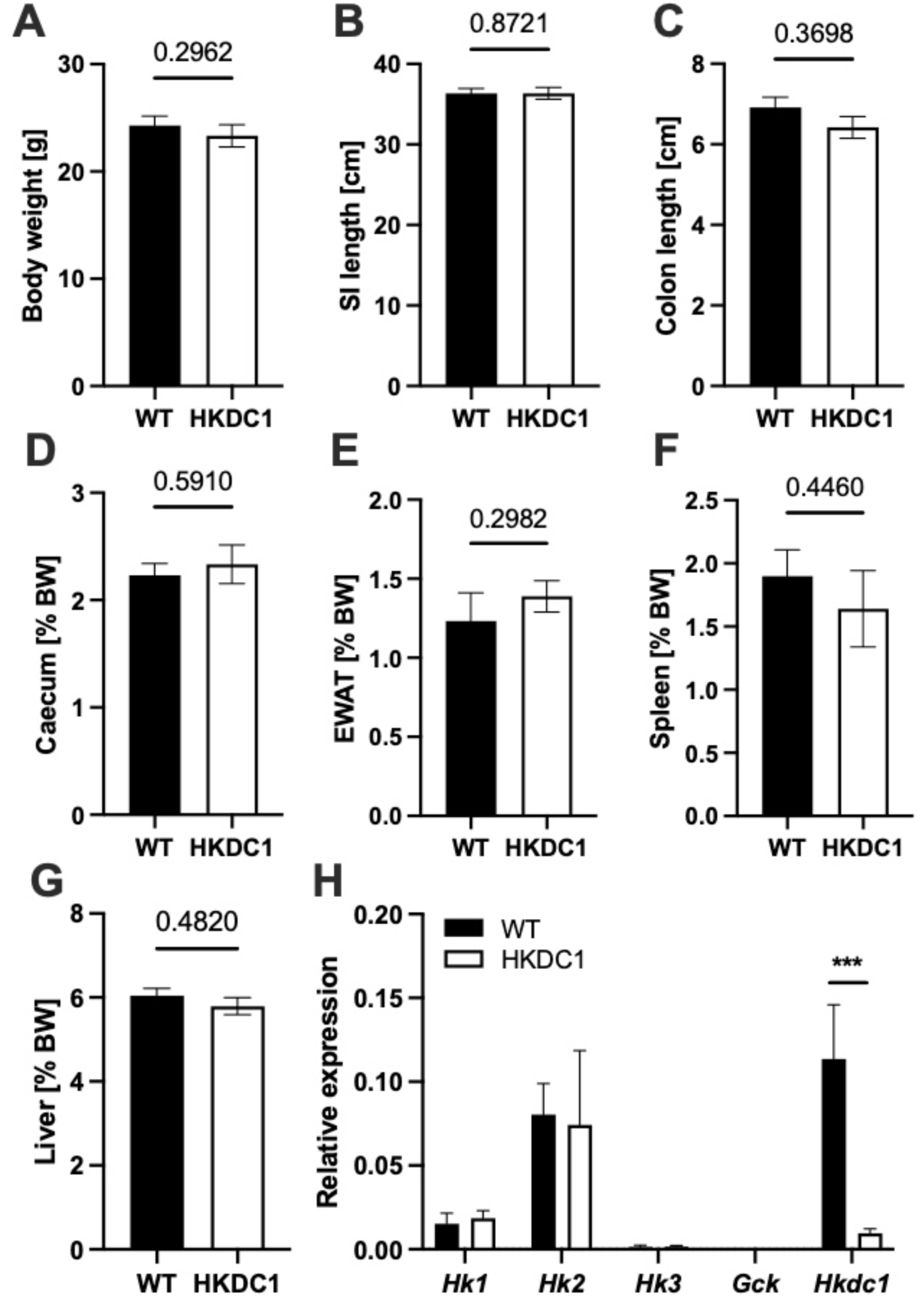
Organ measures did not differ between *Apc*^Min/+^-*Hkdc1*^ΔIEC^ mice and WT controls. (A-H) Organ measures were taken from 20-week old mice *Apc*^Min/+^-*Hkdc1*^ΔIEC^ mice and WT littermate controls. **(A)** Body weight (BW). **(B)** Small intestine (SI) length in cm. **(C)** Colon length in cm. **(D)** Caecum weight as percent of BW. **(E)** Epididymal white adipose tissue (EWAT) as percent of BW. **(F)** Spleen weight as percent of BW. **(G)** Liver weight as percent of BW. **(H)** Expression of all HK-family members in small intestinal tissue of *Apc*^Min/+^-*Hkdc1*^ΔIEC^ mice and WT littermate controls. ***: p < 0.001. All data is shown as mean with SEM.

**Table S1:** HKDC1-interacting proteins identified by immunoprecipitation and used for STRING interaction network. . Proteins were quantified in the LFQ dataset and structural proteins were removed. PSM = peptide spectral match.

| Protein name | Protein ID | Protein coverage | # peptides | # unique peptides | PSM | STRING protein identifier |
| --- | --- | --- | --- | --- | --- | --- |
| ALDH1B1 | Q9CZS1 | 39 | 16 | 16 | 80 | 10090.ENSMUSP00000041260 |
| ARF1 | P84078 | 52 | 5 | 4 | 29 | 10090.ENSMUSP00000126120 |
| ARF4 | P61750 | 42 | 5 | 3 | 13 | 10090.ENSMUSP00000022429 |
| ARF5 | P84084 | 29 | 3 | 1 | 8 | 10090.ENSMUSP00000020717 |
| ATP5A1 | Q03265 | 56 | 24 | 24 | 223 | 10090.ENSMUSP00000026495 |
| ATP5B | P56480 | 56 | 16 | 16 | 124 | 10090.ENSMUSP00000026459 |
| C1RA | Q8CG16 | 28 | 15 | 15 | 44 | 10090.ENSMUSP00000063707 |
| C1S1 | E9Q6C2 | 30 | 14 | 7 | 39 | 10090.ENSMUSP00000125531 |
| CFL1 | P18760 | 60 | 8 | 7 | 54 | 10090.ENSMUSP00000147514 |
| CLCA1 | Q9D7Z6 | 37 | 26 | 26 | 129 | 10090.ENSMUSP00000029919 |
| DBT | P53395 | 71 | 26 | 26 | 161 | 10090.ENSMUSP00000000349 |
| DES | P31001 | 76 | 37 | 32 | 363 | 10090.ENSMUSP00000027409 |
| EIF4A1 | P60843 | 39 | 10 | 10 | 48 | 10090.ENSMUSP00000127034 |
| GAPDH | A0A0A0MQF6 | 62 | 14 | 14 | 144 | 10090.ENSMUSP00000113942 |
| GFPT1 | P47856 | 28 | 13 | 13 | 49 | 10090.ENSMUSP00000109288 |
| HK1 | P17710 | 59 | 53 | 46 | 507 | 10090.ENSMUSP00000111946 |
| HK2 | O08528 | 53 | 41 | 36 | 267 | 10090.ENSMUSP00000000642 |
| HKDC1 | Q91W97 | 64 | 57 | 52 | 524 | 10090.ENSMUSP00000020277 |
| HSPA8 | P63017 | 39 | 19 | 15 | 128 | 10090.ENSMUSP00000015800 |
| LDHA | A0A1B0GSX0 | 26 | 9 | 9 | 71 | 10090.ENSMUSP00000148107 |
| LGALS2 | Q9CQW5 | 64 | 7 | 7 | 32 | 10090.ENSMUSP00000036598 |
| LMNA | P48678 | 59 | 31 | 30 | 140 | 10090.ENSMUSP00000029699 |
| MPTX2 | D3YYJ7 | 37 | 7 | 7 | 28 | 10090.ENSMUSP00000106855 |
| PHB | P67778 | 33 | 5 | 5 | 13 | 10090.ENSMUSP00000119603 |
| PRDX1 | P35700 | 42 | 6 | 4 | 36 | 10090.ENSMUSP00000114159 |
| RACK1 | P68040 | 32 | 7 | 7 | 41 | 10090.ENSMUSP00000020640 |
| SCIN | Q60604 | 71 | 35 | 35 | 286 | 10090.ENSMUSP00000002640 |
| SLC25A4 | P48962 | 40 | 12 | 5 | 112 | 10090.ENSMUSP00000034049 |
| SLC25A5 | P51881 | 42 | 12 | 5 | 150 | 10090.ENSMUSP00000016463 |
| VDAC2 | Q60930 | 54 | 10 | 10 | 59 | 10090.ENSMUSP00000022293 |
| VDAC3 | Q60931 | 31 | 6 | 5 | 26 | 10090.ENSMUSP00000009036 |
| VIL1 | Q62468 | 80 | 55 | 55 | 1241 | 10090.ENSMUSP00000027366 |
| VIM | P20152 | 71 | 33 | 26 | 329 | 10090.ENSMUSP00000028062 |
| ZG16 | Q8K0C5 | 65 | 6 | 6 | 63 | 10090.ENSMUSP00000145876 |

**Table S2:**
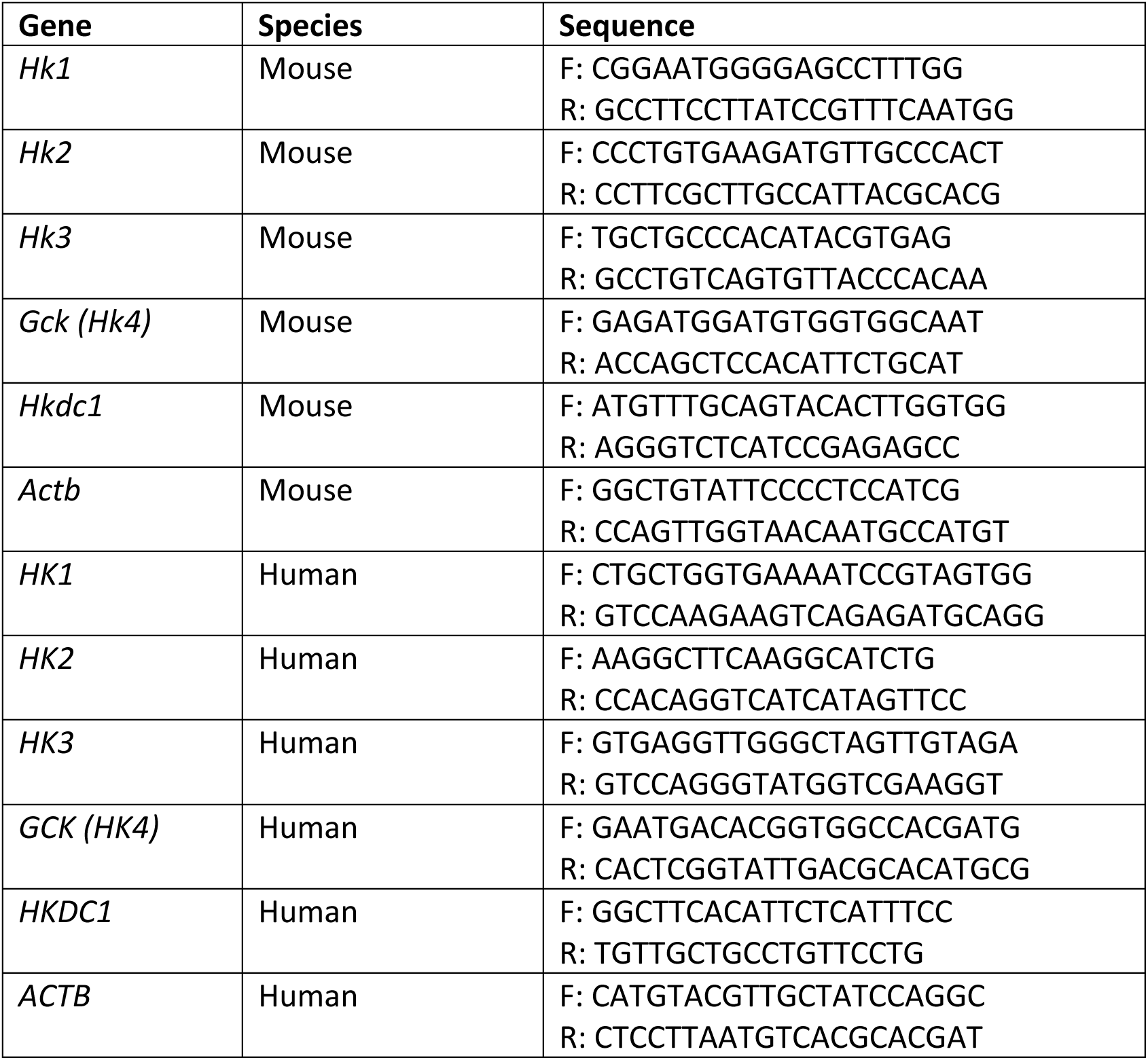
qPCR Primers. Sequences of forward (F) and reverse (R) primers used for RT-PCR analysis.

**Table S3:**
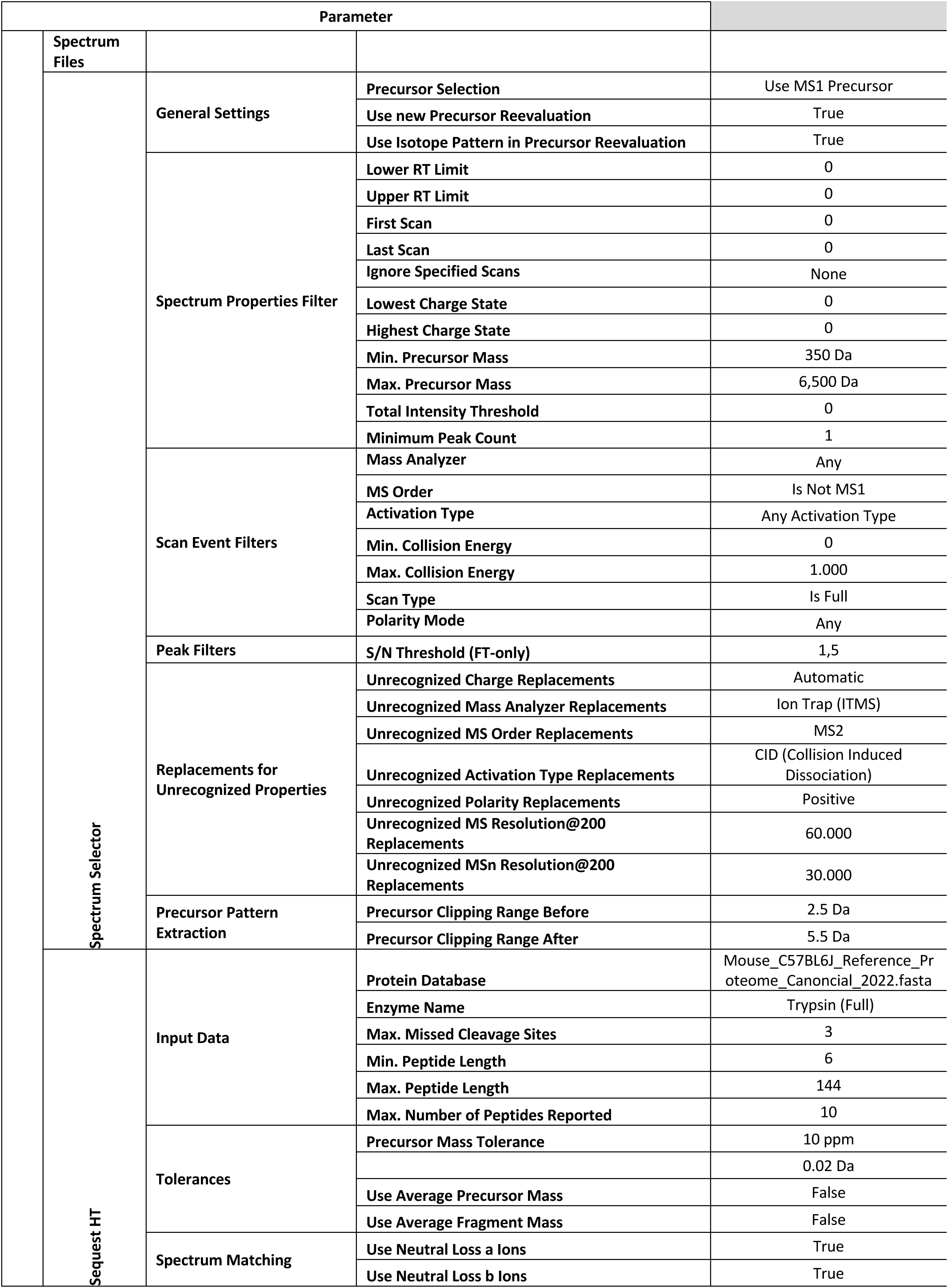
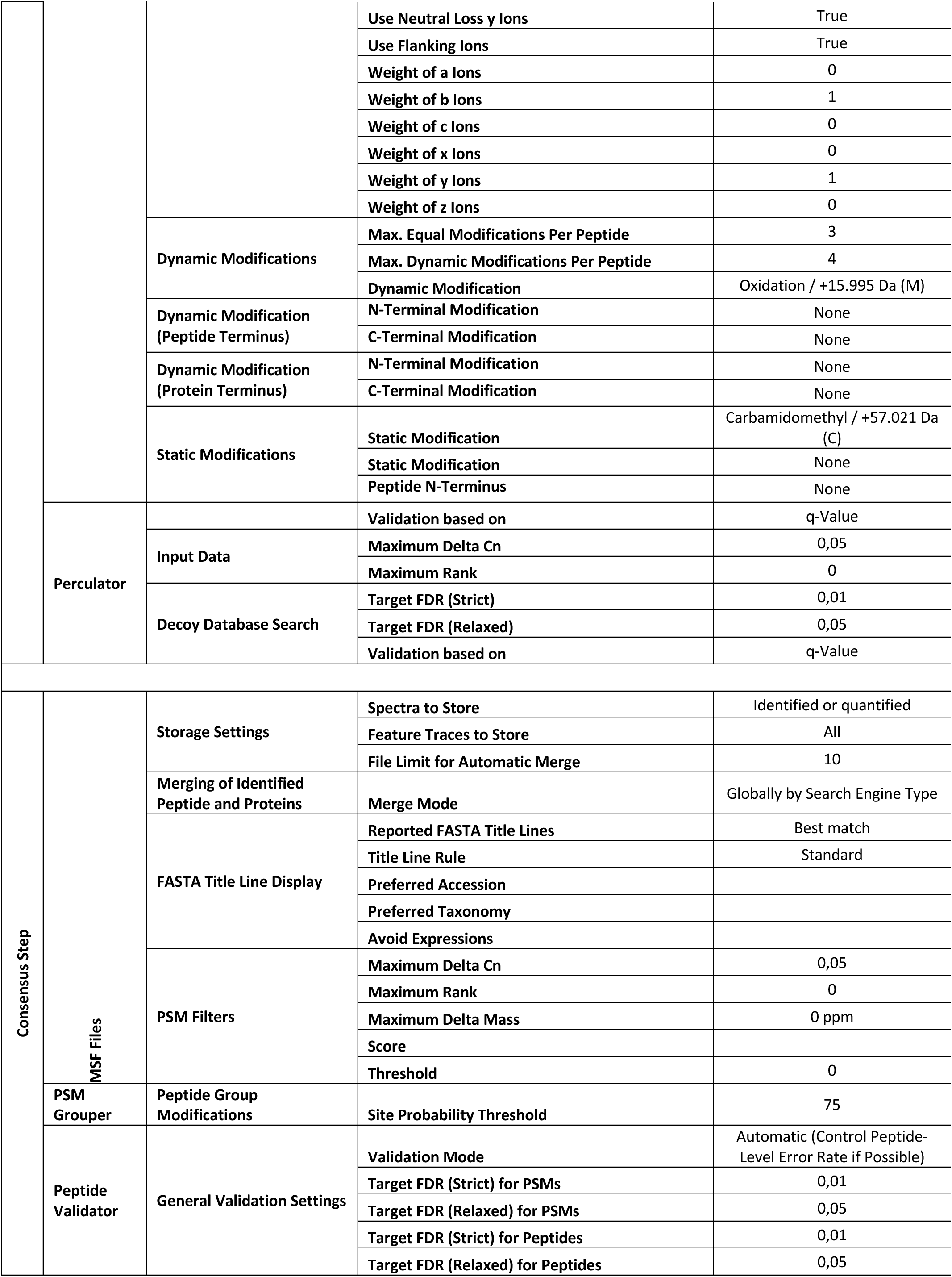
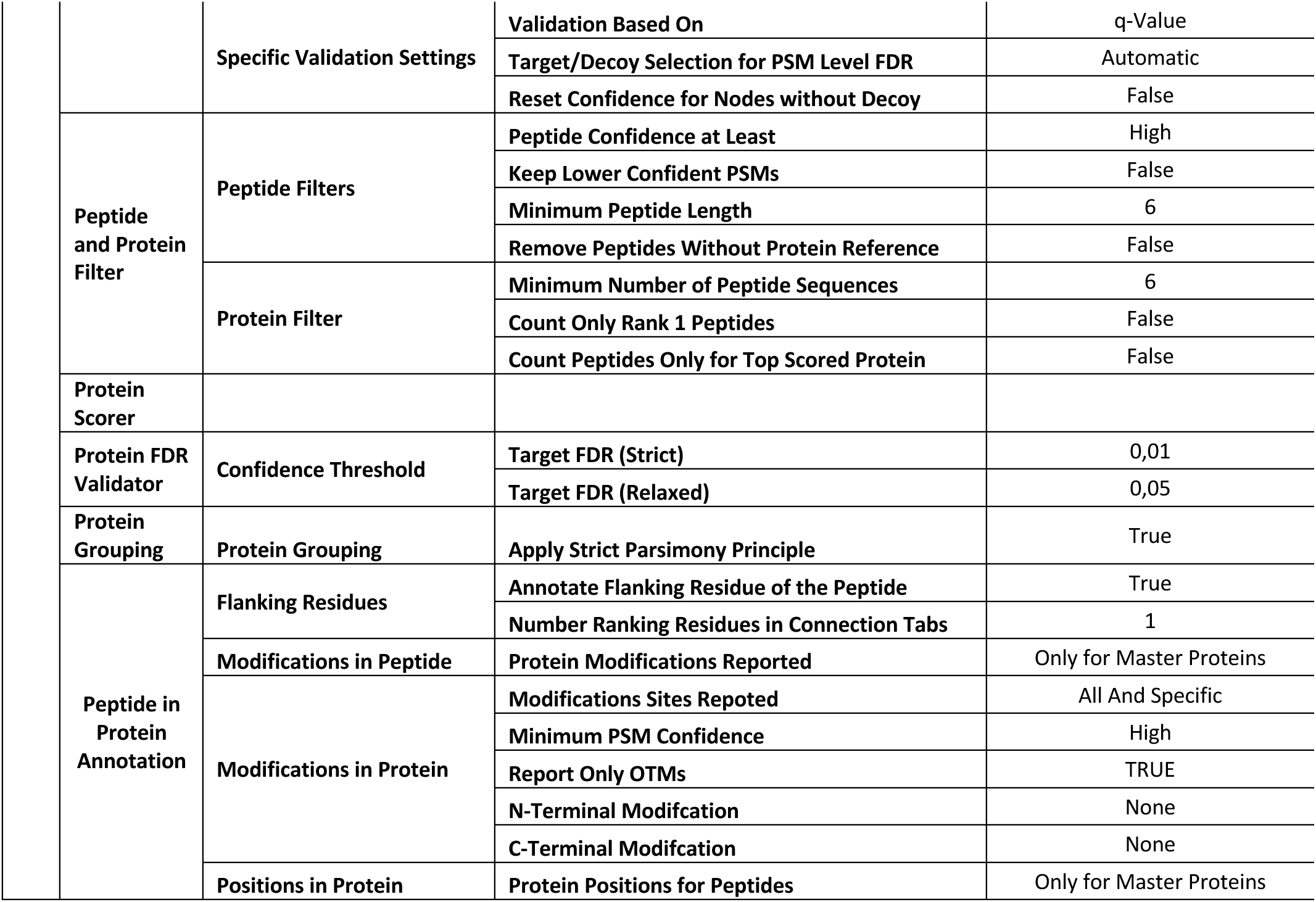
Proteome Discoverer settings. Protein identification parameters used in the Proteome Discoverer (version 2.2.0.388) workflow.

## Abbreviations

ANOVA: Analysis of variance
ANT2: Adenine nucleotide translocator 2
ADP: Adenosine diphosphate
*Apc*^Min/+^: Adenomatous-polyposis-coli multiple intestinal neoplasia
ATP: Adenosine triphosphate
ATP5A1: ATP Synthase Subunit Alpha 1 CRC Colorectal cancer
CRISPR: Clustered regularly interspaced short palindromic repeats DNA deoxyribonucleic acid
Dpi: Days post infection
FACS: Fluorescence activated cell sorting
FDR: False discovery rate
FKPM: Median Fragments Per Kilobase of exon per Million reads
GFPT1: Glutamine-Fructose-6-Phosphate Transaminase 1
GTEx: The Genotype-Tissue Expression
GO: Gene ontology
H&E: Hematoxylin and eosin
HC: Healthy controls
HPA: Human Protein Atlas
HK: Hexokinase
HKDC1: Hexokinase domain containing 1
IEC: Intestinal epithelial cell
IFN-β: Interferon beta
LFQ: Label free quantification
MPTP: Mitochondrial permeability transition pore
NSG: NOD.Cg-Prkdc^SCID^ Il2rg^tm1Wjl^/SzJ
nTPM: normalized protein-coding transcripts per million
PBS: Phosphate-buffered saline
PHB2: Prohibitin 2
RNA: Ribonucleic acid
SEM: Standard error of the mean
SLC25A5: Solute Carrier Family 25 Member 5
TCGA: The Cancer Genome Atlas
TMRM: Tetramethylrhodamin-methylester
TNF: Tumor necrosis factor
TUNEL: Terminal deoxynucleotidyl transferase dUTP nick end labelling
VDAC: Voltage-dependent anion channel
WT: Wildtype

## Declarations

### Ethics approval and consent to participate

All animal experiments were approved by the local animal safety review board of the federal ministry of Schleswig Holstein and conducted according to national and international laws and policies (approval numbers V242-56302/2018(100-11/18) and IX552-65205/2024(24- 4/24)). No human studies were conducted but only data from public databases used.

### Consent for publication

Not applicable

### Availability of data and material

All data is either included in this manuscript or deposited on public databases. The RNA sequencing data are accessible through the European Nucleotide Archive (https://www.ebi.ac.uk/ena) under the accession number PRJEB82610.

### Author contributions

LJ, SW, KS, JG and FS designed research. LJ, SW, KS, KM, JG, FAS, LB, MT, BF and FS performed experiments and analyzed the data. LJ, SW and FS prepared the figures. LJ and FS obtained funding. LJ, SW and FS co-wrote the manuscript with critical input from all authors. All authors read and approved the final manuscript.

## Acknowledgements

The authors thank Sabine Kock, Stefanie Baumgarten, Maren Reffelmann, Vivian Wegner, Tanja Klostermeier, Dorina Ölsner, Meike Hansen, Ronja Möhring and Sophie Reiher for excellent technical assistance.

## References

Ahlemeyer, B., Klumpp, S., & Krieglstein, J. (2002). Release of cytochrome c into the extracellular space contributes to neuronal apoptosis induced by staurosporine. Brain Research, 934(2), 107–116. 10.1016/S0006-8993(02)02365-X

Baik, S. H., Ramanujan, V. K., Becker, C., Fett, S., Underhill, D. M., & Wolf, A. J. (2023). Hexokinase dissociation from mitochondria promotes oligomerization of VDAC that facilitates NLRP3 inflammasome assembly and activation. Science Immunology, 8(84), eade7652. 10.1126/sciimmunol.ade7652

Baines, C. P., & Gutiérrez-Aguilar, M. (2018). The still uncertain identity of the channel- forming unit(s) of the mitochondrial permeability transition pore. Cell Calcium, 73, 121–130. 10.1016/j.ceca.2018.05.003

Baines, C. P., Kaiser, R. A., Purcell, N. H., Blair, N. S., Osinska, H., Hambleton, M. A., Brunskill, E. W., Sayen, M. R., Gottlieb, R. A., Dorn, G. W., Robbins, J., & Molkentin, J. D. (2005). Loss of cyclophilin D reveals a critical role for mitochondrial permeability transition in cell death. Nature, 434(7033), 658–662. 10.1038/nature03434

Carlson, M. (2024). org.Mm.eg.db: Genome wide annotation for Mouse (Version 3.19.1) [R package]. doi:10.18129/B9.bioc.org.Mm.eg.db

Chen, Q., Feng, J., Wu, J., Yu, Z., Zhang, W., Chen, Y., Yao, P., & Zhang, H. (2020). HKDC1 C-terminal based peptides inhibit extranodal natural killer/T-cell lymphoma by modulation of mitochondrial function and EBV suppression. Leukemia, 34(10), 2736– 2748. 10.1038/s41375-020-0801-5

Chen, X., Lv, Y., Sun, Y., Zhang, H., Xie, W., Zhong, L., Chen, Q., Li, M., Li, L., Feng, J., Yao, A., Zhang, Q., Huang, X., Yu, Z., & Yao, P. (2019). PGC1β Regulates Breast Tumor Growth and Metastasis by SREBP1-Mediated HKDC1 Expression. Frontiers in Oncology, 9, 290. 10.3389/fonc.2019.00290

Chen, Y.-J., Hong, W.-F., Liu, M.-L., Guo, X., Yu, Y.-Y., Cui, Y.-H., Liu, T.-S., & Liang, L. (2022). An integrated bioinformatic investigation of mitochondrial solute carrier family 25 (SLC25) in colon cancer followed by preliminary validation of member 5 (SLC25A5) in tumorigenesis. Cell Death & Disease, 13(3), 237. 10.1038/s41419-022-04692-1

Chevrollier, A., Loiseau, D., Reynier, P., & Stepien, G. (2011). Adenine nucleotide translocase 2 is a key mitochondrial protein in cancer metabolism. Biochimica et Biophysica Acta (BBA) - Bioenergetics, 1807(6), 562–567. 10.1016/j.bbabio.2010.10.008

Creed, S., & McKenzie, M. (2019). Measurement of Mitochondrial Membrane Potential with the Fluorescent Dye Tetramethylrhodamine Methyl Ester (TMRM). Methods in Molecular Biology. 10.1007/978-1-4939-9027-6_5

Cui, M., Yamano, K., Yamamoto, K., Yamamoto-Imoto, H., Minami, S., Yamamoto, T., Matsui, S., Kaminishi, T., Shima, T., Ogura, M., Tsuchiya, M., Nishino, K., Layden, B. T., Kato, H., Ogawa, H., Oki, S., Okada, Y., Isaka, Y., Kosako, H., … Nakamura, S. (2024). HKDC1, a target of TFEB, is essential to maintain both mitochondrial and lysosomal homeostasis, preventing cellular senescence. Proceedings of the National Academy of Sciences of the United States of America, 121(2), e2306454120. 10.1073/pnas.2306454120

Czulkies, B. A., Mastroianni, J., Lutz, L., Lang, S., Schwan, C., Schmidt, G., Lassmann, S., Zeiser, R., Aktories, K., & Papatheodorou, P. (2017). Loss of LSR affects epithelial barrier integrity and tumor xenograft growth of CaCo-2 cells. Oncotarget, 8(23), 37009–37022. 10.18632/oncotarget.10425

Dong, L., Lu, D., Chen, R., Lin, Y., Zhu, H., Zhang, Z., Cai, S., Cui, P., Song, G., Rao, D., Yi, X., Wu, Y., Song, N., Liu, F., Zou, Y., Zhang, S., Zhang, X., Wang, X., Qiu, S., … Fan, J. (2022). Proteogenomic characterization identifies clinically relevant subgroups of intrahepatic cholangiocarcinoma. Cancer Cell, 40(1), 70–87.e15. 10.1016/j.ccell.2021.12.006

Franks, L. M., & Hemmings, V. J. (1978). A cell line from an induced carcinoma of mouse rectum. The Journal of Pathology, 124(1), 35–38. 10.1002/path.1711240108

Gong, Y., Qian, Y., Luo, G., Liu, Y., Wang, R., Deng, S., Cheng, H., Jin, K., Ni, Q., Yu, X., Wu, W., & Liu, C. (2021). High GFPT1 expression predicts unfavorable outcomes in patients with resectable pancreatic ductal adenocarcinoma. World Journal of Surgical Oncology, 19(1), 35. 10.1186/s12957-021-02147-z

Guo, J., Ye, F., Xie, W., Zhang, X., Zeng, R., Sheng, W., Mi, Y., & Sheng, X. (2022). The HOXC- AS2/miR-876-5p/HKDC1 axis regulates endometrial cancer progression in a high glucose-related tumor microenvironment. Cancer Science, 113(7), 2297–2310. 10.1111/cas.15384

Hanahan, D. (2022). Hallmarks of Cancer: New Dimensions. Cancer Discovery, 12(1), 31–46. 10.1158/2159-8290.CD-21-1059

Hanahan, D., & Weinberg, R. A. (2000). The Hallmarks of Cancer. Cell, 100(1), 57–70. 10.1016/S0092-8674(00)81683-9

Hanahan, D., & Weinberg, R. A. (2011). Hallmarks of Cancer: The Next Generation. Cell, 144(5), 646–674. 10.1016/j.cell.2011.02.013

Häsler, R., Feng, Z., Bäckdahl, L., Spehlmann, M. E., Franke, A., Teschendorff, A., Rakyan, V. K., Down, T. A., Wilson, G. A., Feber, A., Beck, S., Schreiber, S., & Rosenstiel, P. (2012). A functional methylome map of ulcerative colitis. Genome Research, 22(11), 2130–2137. 10.1101/gr.138347.112

Hayes, M. G., Urbanek, M., Hivert, M.-F., Armstrong, L. L., Morrison, J., Guo, C., Lowe, L. P., Scheftner, D. A., Pluzhnikov, A., Levine, D. M., McHugh, C. P., Ackerman, C. M., Bouchard, L., Brisson, D., Layden, B. T., Mirel, D., Doheny, K. F., Leya, M. V., Lown- Hecht, R. N., … for the HAPO Study Cooperative Research Group. (2013). Identification of *HKDC1* and *BACE2* as Genes Influencing Glycemic Traits During Pregnancy Through Genome-Wide Association Studies. Diabetes, 62(9), 3282–3291. 10.2337/db12-1692

Hu, J., Xia, F., Chen, C., & Zhao, D. (2023a). HKDC1 in Gastric Cancer: A New Diagnostic, Prognostic Biomarker, and Novel Therapeutic Target. Annals of Clinical and Laboratory Science, 53(5), 726–737.

Hu, J., Xia, F., Chen, C., & Zhao, D. (2023b). HKDC1 in Gastric Cancer: A New Diagnostic, Prognostic Biomarker, and Novel Therapeutic Target. Annals of Clinical and Laboratory Science, 53(5), 726–737.

Hughes, C. S., Moggridge, S., Müller, T., Sorensen, P. H., Morin, G. B., & Krijgsveld, J. (2019). Single-pot, solid-phase-enhanced sample preparation for proteomics experiments. Nature Protocols, 14(1), 68–85. 10.1038/s41596-018-0082-x

Hutter, C., & Zenklusen, J. C. (2018). The Cancer Genome Atlas: Creating Lasting Value beyond Its Data. Cell, 173(2), 283–285. 10.1016/j.cell.2018.03.042

Jackson, J. G., O’Donnell, J. C., Krizman, E., & Robinson, M. B. (2015). Displacing hexokinase from mitochondrial voltage-dependent anion channel impairs GLT-1-mediated glutamate uptake but does not disrupt interactions between GLT-1 and mitochondrial proteins. Journal of Neuroscience Research, 93(7), 999–1008. 10.1002/jnr.23533

Jumarie, C., & Malo, C. (1991). Caco-2 cells cultured in serum-free medium as a model for the study of enterocytic differentiation in vitro. Journal of Cellular Physiology, 149(1), 24–33. 10.1002/jcp.1041490105

Käll, L., Storey, J. D., MacCoss, M. J., & Noble, W. S. (2008). Assigning Significance to Peptides Identified by Tandem Mass Spectrometry Using Decoy Databases. Journal of Proteome Research, 7(1), 29–34. 10.1021/pr700600n

Khan, Md. W., Terry, A. R., Priyadarshini, M., Ilievski, V., Farooq, Z., Guzman, G., Cordoba-Chacon, J., Ben-Sahra, I., Wicksteed, B., & Layden, B. T. (2022). The hexokinase “HKDC1” interaction with the mitochondria is essential for liver cancer progression. Cell Death & Disease, 13(7), 660. 10.1038/s41419-022-04999-z

Langlands, A. J., Carroll, T. D., Chen, Y., & Näthke, I. (2018). Chir99021 and Valproic acid reduce the proliferative advantage of Apc mutant cells. Cell Death & Disease, 9(3), 255. 10.1038/s41419-017-0199-9

Li, D., Guan, M., Cao, X., Zha, Z. Q., Zhang, P., Xiang, H., Zhou, Y., Peng, Q., Xu, Z., Lu, L., & Liu, G. (2022). GFPT1 promotes the proliferation of cervical cancer via regulating the ubiquitination and degradation of PTEN. Carcinogenesis, 43(10), 969–979. 10.1093/carcin/bgac073

Li, J., Wang, J., Chen, Y., Yang, L., & Chen, S. (2017). A prognostic 4-gene expression signature for squamous cell lung carcinoma. Journal of Cellular Physiology, 232(12), 3702–3713. 10.1002/jcp.25846

Lian, H., Wang, A., Shen, Y., Wang, Q., Zhou, Z., Zhang, R., Li, K., Liu, C., & Jia, H. (2020). Identification of novel alternative splicing isoform biomarkers and their association with overall survival in colorectal cancer. BMC Gastroenterology, 20(1), 171. 10.1186/s12876-020-01288-x

Lin, J.-S., & Lai, E.-M. (2017). Protein-Protein Interactions: Co-Immunoprecipitation. *Methods in Molecular Biology (Clifton*, N.J.), 1615, 211–219. 10.1007/978-1-4939-7033-9_17

Liu, P., Luo, Y., Wu, H., Han, Y., Wang, S., Liu, R., Wen, S., & Huang, P. (2024). HKDC1 functions as a glucose sensor and promotes metabolic adaptation and cancer growth via interaction with PHB2. Cell Death & Differentiation. 10.1038/s41418-024-01392-5

Locasale, J. W., & Cantley, L. C. (2011). Metabolic Flux and the Regulation of Mammalian Cell Growth. Cell Metabolism, 14(4), 443–451. 10.1016/j.cmet.2011.07.014

Lonsdale, J., Thomas, J., Salvatore, M., Phillips, R., Lo, E., Shad, S., Hasz, R., Walters, G., Garcia, F., Young, N., Foster, B., Moser, M., Karasik, E., Gillard, B., Ramsey, K., Sullivan, S., Bridge, J., Magazine, H., Syron, J., … Moore, H. F. (2013). The Genotype- Tissue Expression (GTEx) project. Nature Genetics, 45(6), 580–585. 10.1038/ng.2653

Love, M. I., Huber, W., & Anders, S. (2014). Moderated estimation of fold change and dispersion for RNA-seq data with DESeq2. Genome Biology, 15(12), 550. 10.1186/s13059-014-0550-8

Ludikhuize, M. C., Meerlo, M., Burgering, B. M. T., & Rodríguez Colman, M. J. (2021). Protocol to profile the bioenergetics of organoids using Seahorse. STAR Protocols, 2(1), 100386. 10.1016/j.xpro.2021.100386

Madison, B. B., Dunbar, L., Qiao, X. T., Braunstein, K., Braunstein, E., & Gumucio, D. L. (2002). Cis Elements of the Villin Gene Control Expression in Restricted Domains of the Vertical (Crypt) and Horizontal (Duodenum, Cecum) Axes of the Intestine. Journal of Biological Chemistry, 277(36), 33275–33283. 10.1074/jbc.M204935200

Mills, E. L., Kelly, B., & O’Neill, L. A. J. (2017). Mitochondria are the powerhouses of immunity. Nature Immunology, 18(5), 488–498. 10.1038/ni.3704

Miyoshi, H., & Stappenbeck, T. S. (2013). In vitro expansion and genetic modification of gastrointestinal stem cells in spheroid culture. Nature Protocols, 8(12), 2471–2482. 10.1038/nprot.2013.153

Moser, A. R., Luongo, C., Gould, K. A., McNeley, M. K., Shoemaker, A. R., & Dove, W. F. (1995). ApcMin: A mouse model for intestinal and mammary tumorigenesis. European Journal of Cancer (Oxford, England: 1990), 31A(7–8), 1061–1064. 10.1016/0959-8049(95)00181-h

Pusec, C. M., De Jesus, A., Khan, M. W., Terry, A. R., Ludvik, A. E., Xu, K., Giancola, N., Pervaiz, H., Daviau Smith, E., Ding, X., Harrison, S., Chandel, N. S., Becker, T. C., Hay, N., Ardehali, H., Cordoba-Chacon, J., & Layden, B. T. (2019). Hepatic HKDC1 Expression Contributes to Liver Metabolism. Endocrinology, 160(2), 313–330. 10.1210/en.2018-00887

Pusec, C. M., Ilievski, V., De Jesus, A., Farooq, Z., Zapater, J. L., Sweis, N., Ismail, H., Khan, M. W., Ardehali, H., Cordoba-Chacon, J., & Layden, B. T. (2023). Liver-specific overexpression of HKDC1 increases hepatocyte size and proliferative capacity. Scientific Reports, 13(1), 8034. 10.1038/s41598-023-33924-3

Quraishi, M. N., Acharjee, A., Beggs, A. D., Horniblow, R., Tselepis, C., Gkoutos, G., Ghosh, S., Rossiter, A. E., Loman, N., van Schaik, W., Withers, D., Walters, J. R. F., Hirschfield, G. M., & Iqbal, T. H. (2020). A Pilot Integrative Analysis of Colonic Gene Expression, Gut Microbiota, and Immune Infiltration in Primary Sclerosing Cholangitis-Inflammatory Bowel Disease: Association of Disease With Bile Acid Pathways. Journal of Crohn’s and Colitis, 14(7), 935–947. 10.1093/ecco-jcc/jjaa021

R Core Team. (2024). R: A Language and Environment for Statistical Computing. R Foundation for Statistical Computing.

Sato, T., Stange, D. E., Ferrante, M., Vries, R. G. J., Van Es, J. H., Van Den Brink, S., Van Houdt, W. J., Pronk, A., Van Gorp, J., Siersema, P. D., & Clevers, H. (2011). Long-term Expansion of Epithelial Organoids From Human Colon, Adenoma, Adenocarcinoma, and Barrett’s Epithelium. Gastroenterology, 141(5), 1762–1772. 10.1053/j.gastro.2011.07.050

Schneikert, J., & Behrens, J. (2007). The canonical Wnt signalling pathway and its APC partner in colon cancer development. Gut, 56(3), 417–425. 10.1136/gut.2006.093310

Siegel, R. L., Miller, K. D., Wagle, N. S., & Jemal, A. (2023). Cancer statistics, 2023. CA: A Cancer Journal for Clinicians, 73(1), 17–48. 10.3322/caac.21763

Siegel, R. L., Wagle, N. S., Cercek, A., Smith, R. A., & Jemal, A. (2023). Colorectal cancer statistics, 2023. CA: A Cancer Journal for Clinicians, 73(3), 233–254. 10.3322/caac.21772

Szklarczyk, D., Kirsch, R., Koutrouli, M., Nastou, K., Mehryary, F., Hachilif, R., Gable, A. L., Fang, T., Doncheva, N. T., Pyysalo, S., Bork, P., Jensen, L. J., & von Mering, C. (2023). The STRING database in 2023: Protein–protein association networks and functional enrichment analyses for any sequenced genome of interest. Nucleic Acids Research, 51(D1), D638–D646. 10.1093/nar/gkac1000

The Cancer Genome Atlas Research Network, Weinstein, J. N., Collisson, E. A., Mills, G. B., Shaw, K. R. M., Ozenberger, B. A., Ellrott, K., Shmulevich, I., Sander, C., & Stuart, J. M. (2013). The Cancer Genome Atlas Pan-Cancer analysis project. Nature Genetics, 45(10), 1113–1120. 10.1038/ng.2764

Uhlén, M., Fagerberg, L., Hallström, B. M., Lindskog, C., Oksvold, P., Mardinoglu, A., Sivertsson, Å., Kampf, C., Sjöstedt, E., Asplund, A., Olsson, I., Edlund, K., Lundberg, E., Navani, S., Szigyarto, C. A.-K., Odeberg, J., Djureinovic, D., Takanen, J. O., Hober, S., … Pontén, F. (2015). Tissue-based map of the human proteome. Science, 347(6220), 1260419. 10.1126/science.1260419

Uronis, J. M., & Threadgill, D. W. (2009). Murine models of colorectal cancer. Mammalian Genome, 20(5), 261–268. 10.1007/s00335-009-9186-5

Van Puyvelde, B., Daled, S., Willems, S., Gabriels, R., Gonzalez De Peredo, A., Chaoui, K., Mouton-Barbosa, E., Bouyssié, D., Boonen, K., Hughes, C. J., Gethings, L. A., Perez- Riverol, Y., Bloomfield, N., Tate, S., Schiltz, O., Martens, L., Deforce, D., & Dhaenens, M. (2022). A comprehensive LFQ benchmark dataset on modern day acquisition strategies in proteomics. Scientific Data, 9(1), 126. 10.1038/s41597-022-01216-6

Wang, M., Chen, Y., Xu, H., Zhan, J., Suo, D., Wang, J., Ma, Y., Guan, X., Li, Y., & Zhu, S. (2023). HKDC1 upregulation promotes glycolysis and disease progression, and confers chemoresistance onto gastric cancer. Cancer Science, 114(4), 1365–1377. 10.1111/cas.15692

Wang, X., Shi, B., Zhao, Y., Lu, Q., Fei, X., Lu, C., Li, C., & Chen, H. (2020). HKDC1 promotes the tumorigenesis and glycolysis in lung adenocarcinoma via regulating AMPK/mTOR signaling pathway. Cancer Cell International, 20(1), 450. 10.1186/s12935-020-01539-7

Warburg, O., Wind, F., & Negelein, E. (1927). The metabolism of tumors in the body. Journal of General Physiology, 8(6), 519–530. 10.1085/jgp.8.6.519

Weiser, M., Simon, J. M., Kochar, B., Tovar, A., Israel, J. W., Robinson, A., Gipson, G. R., Schaner, M. S., Herfarth, H. H., Sartor, R. B., McGovern, D. P. B., Rahbar, R., Sadiq, T. S., Koruda, M. J., Furey, T. S., & Sheikh, S. Z. (2018). Molecular classification of Crohn’s disease reveals two clinically relevant subtypes. Gut, 67(1), 36–42. 10.1136/gutjnl-2016-312518

Wu, T., Hu, E., Xu, S., Chen, M., Guo, P., Dai, Z., Feng, T., Zhou, L., Tang, W., Zhan, L., Fu, X., Liu, S., Bo, X., & Yu, G. (2021). clusterProfiler 4.0: A universal enrichment tool for interpreting omics data. The Innovation, 2(3), 100141. 10.1016/j.xinn.2021.100141

Yu, C., Bao, T., Jin, L., Lu, J., & Feng, J. (2023). HKDC1 Silencing Inhibits Proliferation and Glycolysis of Gastric Cancer Cells. Journal of Oncology, 2023, 1–15. 10.1155/2023/3876342

Zapater, J. L., Lednovich, K. R., Khan, Md. W., Pusec, C. M., & Layden, B. T. (2022). Hexokinase domain-containing protein-1 in metabolic diseases and beyond. Trends in Endocrinology & Metabolism, 33(1), 72–84. 10.1016/j.tem.2021.10.006

Zhang, G., Zhong, J., Lin, L., & Liu, Z. (2022). Loss of ATP5A1 enhances proliferation and predicts poor prognosis of colon adenocarcinoma. Pathology - Research and Practice, 230, 153679. 10.1016/j.prp.2021.153679

Zhang, Y., Wang, M., Ye, L., Shen, S., Zhang, Y., Qian, X., Zhang, T., Yuan, M., Ye, Z., Cai, J., Meng, X., Qiu, S., Liu, S., Liu, R., Jia, W., Yang, X., Zhang, H., Zhong, X., & Gao, P. (2024). HKDC1 promotes tumor immune evasion in hepatocellular carcinoma by coupling cytoskeleton to STAT1 activation and PD-L1 expression. Nature Communications, 15(1), 1314. 10.1038/s41467-024-45712-2

Zhao, P., Yuan, F., Xu, L., Jin, Z., Liu, Y., Su, J., Yuan, L., Peng, L., Wang, C., & Zhang, G. (2023). HKDC1 reprograms lipid metabolism to enhance gastric cancer metastasis and cisplatin resistance via forming a ribonucleoprotein complex. Cancer Letters, 569, 216305. 10.1016/j.canlet.2023.216305

